# Organ-wide and ploidy-dependent regulations both contribute to cell size determination: evidence from a computational model of tomato fruit

**DOI:** 10.1101/450916

**Authors:** Valentina Baldazzi, Pierre Valsesia, Michel Génard, Nadia Bertin

## Abstract

The development of a new organ is the result of coordinated events of cell division and expansion, in strong interaction with each other. This paper presents a dynamic model of tomato fruit development that includes cell division, endoreduplication and expansion processes. The model is used to investigate the potential interaction among these developmental processes, in the perspective of the neo-cellular theory. In particular, different control schemes (either cell-autonomous or organ-controlled) are tested and compared to experimental data related to two contrasted genotypes. The model shows that a pure cell-autonomous control fails to reproduce the observed cell size distribution, and an organ-wide control is required in order to get realistic cell size variations. The model also supports the role of endoreduplication as an important determinant of final cell size and suggests that a direct effect of endoreduplication on cell expansion is needed in order to obtain a significant correlation between size and ploidy, as observed in real data.

## INTRODUCTION

Understanding the mechanisms underpinning fruit development from its early stages is of primary importance for biology and agronomy. Indeed, early stages are highly sensitive to biotic and abiotic stresses, with important consequences on fruit set and yield. The development of a new organ is the result of coordinated events of cell division and expansion. Fruit growth starts immediately after pollination with intensive cell division. As development proceeds, the proliferative activity of cells progressively slows down giving way to a phase of pure cell enlargement during fruit growth and ripening. In many species, including tomato, the transition from cell division to expansion phases is accompanied by repeated DNA duplications without mitosis, a process called endoreduplication. The exact role of endoreduplication is still unclear. A significant correlation between cell ploidy (i.e number of DNA copies) and cell size has been observed in different species, including tomato fruit, suggesting a possible role of endoreduplication into the control of organ growth Breuer et al., 2010; Chevalier et al., 2011; Cheniclet, 2005). Several studies however showed that, under specific conditions, the two processes can be uncoupled to some extent, so that ploidy is not the only determinant of cell size (Bertin, 2005; Cookson *et al*., 2006).

Understanding the way cell division, endoreduplication and expansion processes interact is crucial to predict the emergence of important morphological traits (fruit size, mass, shape and texture) and their dependence on environmental and genetic factors. Historically, a big debate has opposed two contrasting views, the cellular vs the organismal theory, that set the control of organ growth at the level of the individual cell or of the whole tissue, respectively (reviewed in Beemster et al., 2003; Fleming, 2006; John and Qi, 2008). In the recent years, a consensus view, the neo-cellular theory, has eventually emerged. Accordingly, although cells are the units of plant morphology, their behavior (division, expansion) is not autonomous, but coordinated at the organ level by cell-to-cell communication mechanisms (Beemster et al., 2003; Sablowski and Carnier Dornelas, 2014; Tsukaya, 2003). The existence of non-cell autonomous control of organ development has been demonstrated in Arabidopsis leaf (Kawade et al., 2010) but the underlying modes of action remain unclear and often species-or organ-specific (Ferjani et al., 2007; Han et al., 2014; Horiguchi and Tsukaya, 2011; Norman et al., 2011; Okello et al., 2015).

Computational models offer a unique tool to express and test biological hypotheses, in a well-defined and controlled manner. Not surprisingly, indeed, computational modeling has been largely used to investigate the relationships between organ development and the underlying cellular processes. Many works have addressed the question of organogenesis, relating local morphogenetic rules and cell mechanical properties with the emerging patterns near the meristem (Boudon et al., 2015; Dupuy et al., 2010; Kuchen et al., 2012; Löfke et al., 2015; Lucas et al., 2013; Robinson et al., 2011; von Wangenheim et al., 2016). At the tissue scale, a few models have addressed the issue of cell size variance based on observed kinematic patterns of cell division or growth rates, with a particular attention to the intrinsic stochasticity of cell-cycle related processes (Asl et al., 2011; Kawade and Tsukaya, 2017; Roeder et al., 2010). In most of these models, cell expansion is simply described via an average growth rate, possibly modulated by the ploidy level of the cell, without any reference to the underlying molecular mechanisms or to the environmental conditions.

To our knowledge, very few attempts have been made to explicitly model the interaction among cell division, expansion and endoreduplication and at the scale of organ development. In Fanwoua et al., 2013 a model of tomato fruit development has been developed that integrates cell division, expansion and endoreduplication processes based on a set of biologically-inspired rules. The fruit is described by a set of *q* classes of cells with the same age, ploidy and mass. Within each class, cell division and endoreduplication are described as discrete events that take place within a well-defined window of time, whenever a specific mass-to-ploidy threshold is reached. Cell growth in dry mass is modeled following a source-sink approach as a function of thermal time, cell’s ploidy and external resources. The model is able to qualitatively capture the effect of environmental conditions (temperature, fruit load) on the final fruit dry mass, but hypotheses and parameters are hard to validate as comparison to experimental data is lacking. Moreover, the water content of the cell is not considered preventing the analysis of cell volumes.

Baldazzi and coworkers recently developed an integrated model of tomato fruit development which explicitly accounts for the dynamics of cell proliferation as well as for the mechanisms of cell expansion, in both dry and fresh masses, based on biophysical and thermodynamical principles (Baldazzi et al., 2012, 2013). Here, a new version of this model, that includes cell endoreduplication is proposed. The model was used to investigate different hypotheses concerning the regulation and the interaction among cellular processes, with special attention to 1) the importance of an organ-wide regulation on cell growth and 2) the potential effect of endoreduplication on cell expansion.

We focus on wild-type organ development and we analyze the effect of organ-wide or cell ploidy-dependent regulation onto the dynamics of cell expansion. To this aim, different control schemes (either cell-autonomous or organ-controlled, with our without ploidy effect on cell expansion) were tested *in silico* by means of specific model variants. Simulation results were analyzed and compared to cell size distributions observed in the fruit pericarp of two contrasted genotypes, a cherry and a large-fruited tomato variety.

The model shows that a pure cell-autonomous control cannot reproduce the experimental cell size distribution, and organ-wide and ploidy-dependent controls are required in order to get realistic cell sizes. In particular, a direct effect of endoreduplication on cell expansion was needed in order to obtain a significant correlation between size and ploidy, as observed in real data.

## MATERIALS AND METHODS

### Experimental data

Two datasets were collected from two glasshouse experiments performed at INRA Avignon (south of France) in 2004 and 2007 on large-fruited (cv Levovil) and cherry (cv. Cervil) tomato genotypes of *Solanum lycospersicum* L. The 2004 experiment fruits were collected from April to May (planting in February) whereas in the 2007 experiment fruits were sampled from October to December (planting in August). Plants were grown according to standard cultural practices. Trusses were pruned in order to homogenize truss size along the stem within each genotype. The maximum number of flowers left on each inflorescence was 12 for Cervil and 6 for Levovil. Flowers were pollinated by bumblebees. Air temperature and humidity were recorded hourly in each experiment and input in the model as external signals.

In both experiments, flower buds and fruits were sampled at different time points relative to the time of flower anthesis (full-flower opening). Fruit fresh and dry mass and pericarp fresh mass were systematically measured at all times points. Pericarp dry mass was estimated assuming a dry mass content equivalent to that of the whole fruit.

In 2004, half of the fruit pericarps were then analyzed by flow cytometry and the other half were used for the determination of cell number. The number of pericarp cells was measured after tissue dissociation according to a method adapted from that of Bünger-Kibler and Bangerth, 1982 and detailed in Bertin et al., 2003. Cells were counted in aliquots of the cell suspension under an optical microscope, using Fuchs-Rosenthal chambers or Bürker chambers for the large and small fruits, respectively. Six to 8 aliquots per fruits were observed and the whole pericarp cell number was calculated according to dilution and observation volumes. The ploidy was measured in the pericarp tissue, as described in Bertin et al., 2007. The average value of three measurements per fruit (when allowed for by the fruit’s size), was included in the analysis.

In the 2007 experiment, the dynamics of cell number (but not endoreduplication) was measured following the same method as in the 2004 experiment. In addition, cell size distribution (smallest and largest radii and 2D-surface) distributions were measured with ImageJ software (imagej.nih.gov/ij/) in the cell suspension aliquots. About 20 to 25 cells per fruit pericarp were measured randomly, on several fruits. Cell size distribution were measured on ripe fruits at about 43 days after anthesis (DAA) for Cervil and 60 DAA for Levovil in the considered growing conditions.

### Model description

The model is composed of two interacting modules, both issued from previously published models (Bertin et al., 2007; Fishman and Génard, 1998; Liu et al., 2007). The fruit is described as a collection of cell populations, each one having a specific age, ploidy and volume, which evolve and grow over time during fruit development. Two cell classes are defined: the proliferating cells and the expanding – endoreduplicating cells. The division-endoreduplication module governs the evolution of the number of cells in each classes, their age (initiation date) and ploidy level, based on genotype-specific parameters (Bertin et al., 2007). At each mitotic cycle, a fraction of proliferating cells proceeds through division whereas the remaining ones enter the expansion phase: a new group of expanding cells is created, together with an array of sub-classes of possible ploidy levels *p*. At initialization of the group, all expanding cells are put into the 4C level.

It is assumed that the onset of endoreduplication coincides with the beginning of the expansion phase. As the endocycles proceed, in each group of expanding cells, a fraction σ of the cells increases its ploidy level *p* by a factor 2 and the distribution of cells across the different ploidy levels is updated.

At any time, the mass (both fresh and dry component) of expanding cells is computed by a biophysical expansion module according to cell’s characteristics (age, ploidy) and depending on available resources and environmental conditions (Fishman and Génard, 1998; Liu et al., 2007). Briefly, cell expansion is described by iteratively solving the Lockhart equation relating the rate of volume increase to the cell’s internal pressure and cell’s mechanical properties (Lockhart, 1965). Flows of water and solutes across the membrane are described by thermodynamic equations and depend on environmental conditions. The relative importance of each transport process may vary along fruit developmental stages, depending on specific developmental control. A full description of the model and its equations can be found in the section S2 of the Supplemental Material.

The model assumes that all cells have equal access to external resources, independently from the number of cells (no competition). All the parameters of the division-endoreduplication module are considered to be independent from environmental conditions for the time being.

### Model initialisation and input

The model starts at the end of the pure division phase, when the proliferative activity of the cells declines and the expansion phase begins (Baldazzi et al., 2013). For Cervil genotype this corresponds to approximatively 8 days before anthesis and to 3 days before anthesis for Levovil genotype (Bertin et al., 2007). The initial number of cells for the 2007 experiment, *n0*, was estimated to 3.3e3 for the cherry tomato (Cervil) and 4.6e4 for the large-fruited (Levovil) genotype based on a few measurements. At the beginning of the simulation, all cells are supposed proliferating with a ploidy level of 2C (transient ploidy of 4C during cell cycle is not considered here). Proliferating cells are supposed to have a constant cell mass, *m0*, as often observed in meristematic cells (homogeneity in cell size) (Sablowski and Carnier Dornelas, 2014; Serrano-Mislata et al., 2015).

The initial mass of the fruit is therefore Mf(0)*=n0*\**m0=n0*\*(*w0* +*s0*), where *w0* and *s0* are initial cell water and dry mass, respectively. At any time, cells leaving the proliferative phase start to grow, from an initial mass 2\**m0* and a ploidy level of *4C*, according to the expansion model.

Cell expansion depends on environmental conditions and resources provided by the mother plant. The phloem sugar concentration is assumed to vary daily between 0.15 and 0.35 M whereas stem water potential oscillates between −0.05 and −0.6 MPa i.e. typical pre-dawn and minimal stem water potential measured for the studied genotypes. Temperature and humidity are provided directly by real-time recording of greenhouse climatic conditions.

### Choice of the model variants: control of cell expansion capabilities

In the integrated model, a number of time-dependent functions account for developmental regulations of cell metabolism and physical properties during the expansion phase (Baldazzi et al. 2013, Liu et al.2007). Two characteristic time-scales are recognizable in the model: the *cell age*, i.e. the time spent since an individual cell has left the proliferative phase, and *organ age* i.e. the time spent since the beginning of the simulation (Figure 1). Depending on the settings of the corresponding time-dependent functions, different cellular processes may be put under cell-autonomous or non-cell autonomous control (hereafter indicated as organ-wide control), allowing for an *in silico* exploration of alternative control hypotheses in the perspective of the cellular and organismal theories. Moreover, a direct effect of cell DNA content onto cell expansion capabilities may be tested according to biological evidences (Chevalier et al., 2011; Edgar et al., 2014; Sugimoto-Shirasu and Roberts, 2003).

**Figure 1:**
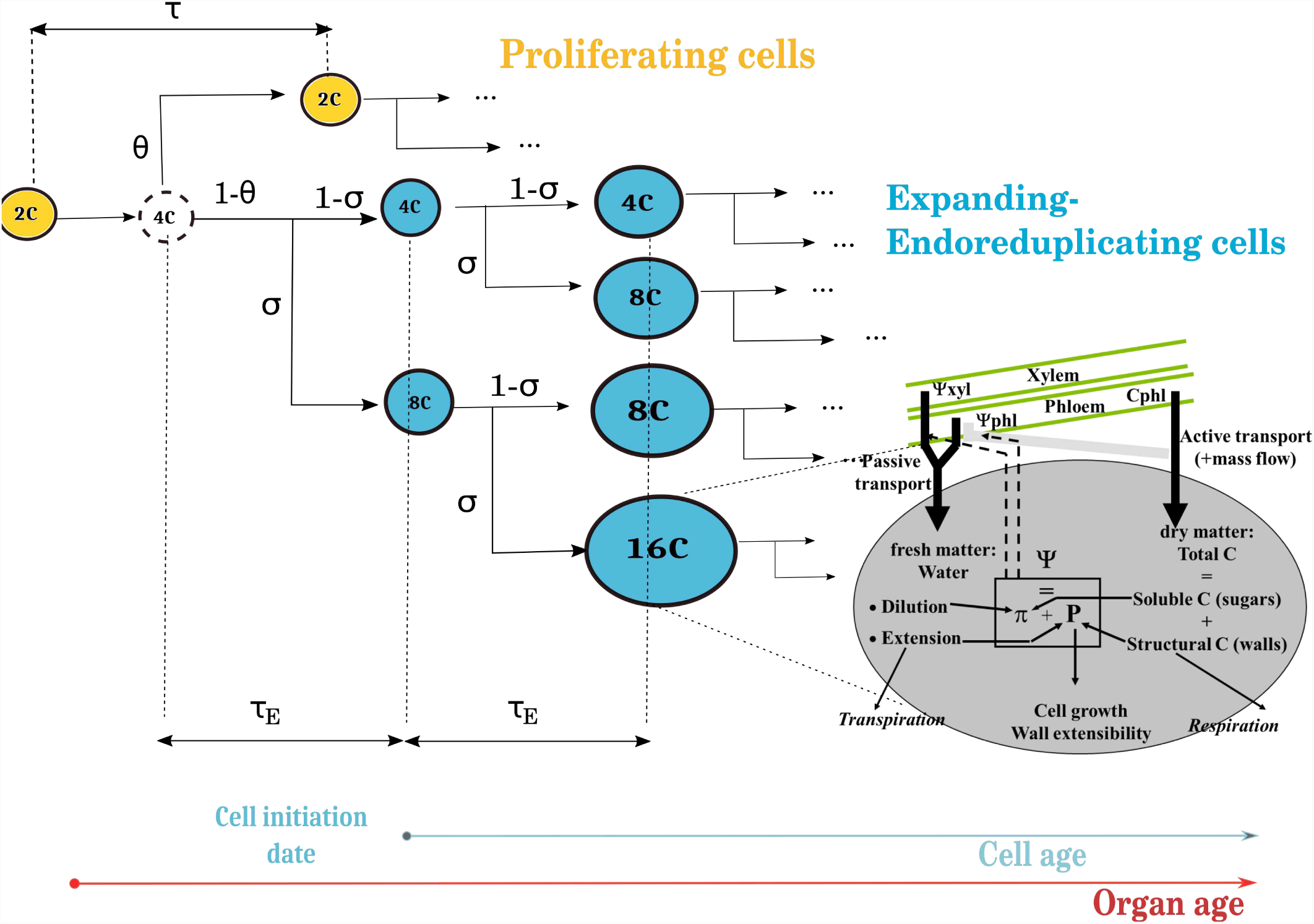
Scheme of the integrated model. The fruit is described as a collection of cell populations, each one having a specific age, ploidy and volume. Cells can be either proliferating or expanding-endoreduplicating. The number of cells in each class is predicted by the division-endoreduplication module, assuming a progressive decline of cells’ proliferating activity. Expanding cells grow according to the expansion module which provides a biophysical description of the main processes involved in carbon and water accumulation. It is assumed that the onset of endoreduplication coincides with the beginning of the expansion phase. Two timescales are recognizable in the model: the organ age i.e. the time since the beginning of the simulation, and the cell age i.e. the time since the cell left the mitotic cycle and entered the expansion-endoreduplication phase. Depending on the model version, cell expansion may be modulated by organ age (organ-wide control) and/or by cell ploidy.

As a default all cellular processes are supposed to depend on cell age (cell-autonomous control) with the only exception of cell transpiration which is computed at the organ scale, on the basis of fruit external surface and skin conductance, and then distributed back to individual cells, proportionally to their relative water content (see section S2).

Based on literature information and on preliminary tests (Baldazzi et al., 2013, 2017) the switch between symplastic and apoplastic transport, σ_*p*_ has been selected as the candidate process for an organ-wide control. Indeed, intercellular movement of macromolecules across plasmodesmata has been shown to be restricted by organ age in tobacco leaves (Crawford and Zambryski, 2001; Zambryski, 2004) and it is known to be important for cell-to-cell communication (Han et al., 2013).

The exact mechanisms by which cell DNA content may affect cell expansion remain currently unknown. Based on literature information and common sense, three distinct mechanisms of action of endoreduplication on cell expansion were hypothesized.

1. Endoreduplication has been often associated to an elevated protein synthesis and transcriptional activity (Chevalier et al., 2014) suggesting a general activation of the nuclear and metabolic machinery of the cell to sustain cell growth (Sugimoto-Shirasu and Roberts, 2003). Following these insights, a first hypothesis assumes an effect of endoreduplication on cell expansion as a ploidy-dependent maximal import rate for carbon uptake. For sake of simplicity, the relation was supposed to be linear in the number of endocycles. The corresponding equation, as a function of the cell DNA content (DNAc, being 2 for dividing cells, 4 to 512 for endoreduplicating cells), was

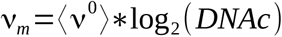

where ⟨ ν^0^⟩ is the average C uptake activity per unit mass.
2. Assuming that cell shape remains the same with increasing ploidy, endoreduplicating cells are characterized by a reduced surface-to-volume ratio with respect to 2C cells (Schoenfelder and Fox, 2015). As a consequence, it is tempting to suppose that one possible advantage of a high ploidy level may reside in a reduction of carbon demand for cell wall and structural units (Barow, 2006; Pirrello et al., 2018). We speculated that such an economy may impact cell expansion capabilities in two ways. First, the metabolic machinery could be redirected towards the synthesis of soluble components, thus contributing to the increase of cell’s internal pressure and consequent volume expansion. In the model, the *ssrat* fraction of soluble compound within the cell is developmentally regulated by the age *t* of the cell (Baldazzi et al. 2013) as

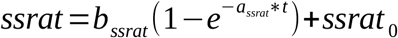 In the presence of a ploidy effect, the final bssrat value was further increased as

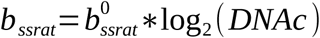
3. Alternatively, “exceeding” carbon may be used to increase the rate of cell wall synthesis or related proteins, possibly resulting in a increase of cell wall plasticity as shown in other systems (Proseus and Boyer, 2006, Jégu *et al*., 2013). In the original expansion model on tomato (Liu et al., 2007) cell wall extensibility Phi declines during cell maturation (Proseus et al., 1999) as

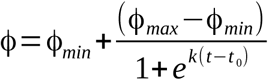 In the presence of a ploidy effect, the maximal cell wall extensibility was increased as

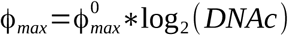 The individual and combined effects of organ-wide and ploidy-dependent control on cell expansion were investigated and compared to a full cell-autonomous model. A total of 10 model variants have been tested for each genotype, following the experimental design shown in Table 1.

**Table 1:** Experimental design showing the characteristics of the 10 model versions tested in the paper.

| Model Variant | ORGAN CONTROL | ENDO EFFECT |  |  |
| --- | --- | --- | --- | --- |
|  | symplastic transport | Active C uptake | Carbon allocation | Wall plasticity |
| M0 |  |  |  |  |
| M1 | ✓ |  |  |  |
| M2 | ✓ | ✓ |  |  |
| M3 | ✓ |  | ✓ |  |
| M4 | ✓ |  |  | ✓ |
| M5 |  | ✓ |  |  |
| M6 |  |  | ✓ |  |
| M7 |  |  |  | ✓ |
| M23 | ✓ | ✓ | ✓ |  |
| M24 | ✓ | ✓ |  | ✓ |

### Model calibration

Calibration has been performed using genetic algorithm under R software (library ‘genalg’). Due to data limitations, a three-steps procedure has been used for each tomato genotype.

First, the division-endoreduplication module (7 parameters) was calibrated on data from the 2004 experiment by comparing measured and simulated values of the total pericarp cell number and the proportion of cells in different ploidy classes, all along fruit development. In particular, this allowed the estimation of the average duration of the endocycle (τ_*E*_) and the proportion (σ) of the cells performing a new endoreduplication round every time τ_*E*_.

The best fitting values of *σ* and τ_*E*_ were selected and kept fixed for the second phase of the calibration, assuming they depend little on environmental conditions (Bertin, 2005). The dynamics of cell division (5 parameters) was then re-estimated on cell number measured in the 2007 experiment, in order to account for environmental regulations of the mitotic cell cycle (see Supplement Information, section S3.1). The best fitting parameters were selected and used for the last calibration step.

The expansion module was calibrated on the evolution of pericarp fresh and dry mass from the 2007 experiment, for which cell size distribution were measured. Six parameters have been selected for calibration based on a previous sensitivity analysis (Constantinescu et al., 2016), whereas the others have been fixed to the original values (Baldazzi et al., 2013; Fishman and Génard, 1998; Liu et al., 2007). An additional parameter was estimated for model variants M3 to M24 in order to correctly evaluate the strength of the ploidy-dependent control (see section S3 for more information).

Due to their different structures, the expansion module was calibrated independently for each model variant. The quality of model adjustment was evaluated using a Normalized Root Mean Square Error (NRMSE):

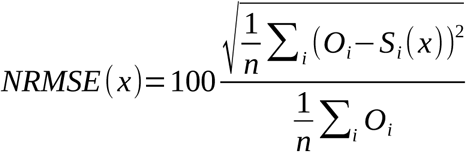

where O_i_ and S_i_ are respectively, the observed and simulated values of pericarp fresh and dry masses, and *n* is the number of observations. *x*={*x*_1_, *x*_2_… *x_p_*} is parameter set of the evaluated solution. The smaller the NRMSE the better the goodness-of-fit is. A NRMSE < 20% is generally considered good, fair if 20% < NRMSE <30% and poor otherwise.

Three to five estimations were performed for each model variant and genotype.

### Solution selection and model comparison

For each calibration solution, the corresponding cell size distribution at fruit maturity (i.e. 43 DAA for Cervil, 60 DAA for Levovil) was reconstructed and compared to the measured data.

A *semi-quantitive* comparison approach has been used due to the limited experimental information available: the general distribution characteristics (shape, positioning and dispersion) have been characterized rather then a perfect fit. To this aim, 8 descriptive statistical indicators *m(i)* have been computed for each solution and compared to those derived from real-data distribution, namely:

- skewness and kurtosis (distribution’s shape)
- mean and median cell size (positioning)
- standard deviation (SD) and median absolute deviation (MAD) (data dispersion)
- maximal and minimal cell size (data dispersion)

Confidence interval (CI, 95%) for the experimental distribution indicators were estimated using a Boostrap approach on 10000 samples.

Based on this scores, the distance between the predicted and the observed distribution has been quantified as the euclidean distance between each indicator *m(i)* and its corresponding measured value, weighted by the amplitude of the confidence interval (DeltaCI) of the indicator itself:

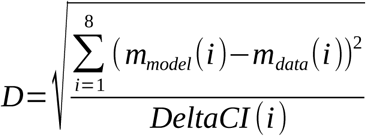

For each model variant, the selection of the best calibration solution has been performed based on a compromise between quality of the fit at the whole fruit scale (as measured by the total NRMSE) and quality of the corresponding cell size distribution (as measured by D, see section S3.2). Estimated parameters for the retained solution are reported in tables S3 and S4.

In order to compare the distributions issued from the different models, a *principal component analysis* (PCA) was performed on the 8 descriptors of cell distribution arising from each model estimation. The ade4 library of R software was used for this purpose (R development Core Team 2006).

## RESULTS

### A characteristic right-tailed distribution of cell areas

The distribution of cell sizes at a given stage of fruit development directly depends on the particular cell division and expansion patterns followed by the organ up to the considered time. Any change in the cell division or expansion rate will have a consequence on the shape and position of the resulting distribution. For both tomato genotypes considered in this study, the distribution of pericarp cell areas at mature stage shows a typical right-tailed shape (Figure 2), compatible with a Weibull or a Gamma distribution (see section S1). The observed cell sizes span up to two orders of magnitude, with cell areas (cross section) ranging from 0.004 to 0.08 mm^2^, for Cervil genotype, and from 0.005 to 0.28 mm2 for Levovil (Table 3 and 4). The average cell area is calculated to be 0.026 mm2 for the cherry tomato and 0.074 mm2 for the large-fruited genotype, values in agreement with data from other tomato varieties (Bertin, 2005; Renaudin et al., 2017). Data dispersion is higher for the large-fruited genotype, but the shape of the distribution, as measured by its skewness and kurtosis values, is pretty similar for both tomato varieties.

**Figure 2:**
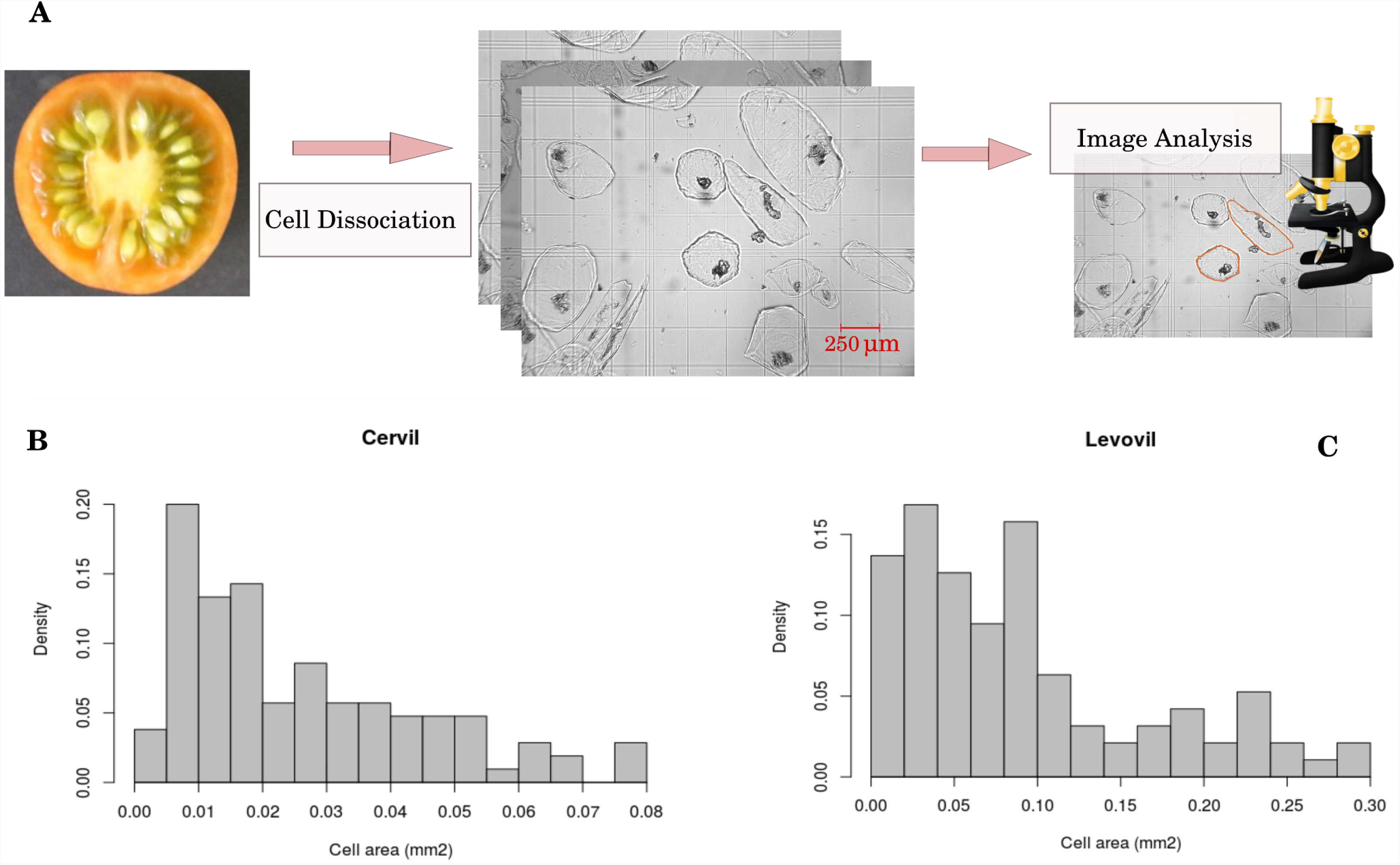
*Top, panel A:* schematic representation of our experimental procedure for cell size estimation. About half of the fruit pericarp underwent tissue dissociation. Cells were divided in different aliquots and spotted onto clean glass slide, before microscope imagining. Images were collected with a digital camera and analysed using ImageJ software. *Bottom*: Measured cell size distribution at fruit maturity *B*: Cervil genotype. *C*: Levovil genotype.

In the following, the effect of specific control mechanisms on the resulting cell area distribution is analysed in details, based on the results obtained for the selected solution (see M&M section, ‘Solution Selection’). The corresponding statistical descriptors are reported in Table 2 and 3, for Cervil and Levovil genotype respectively. Note that predicted minimal cell sizes for Levovil genotype are systematically lower than experimental measurements and correspond to the size of proliferating cells (assumed constant in the present version of the model).

**Table 2:**
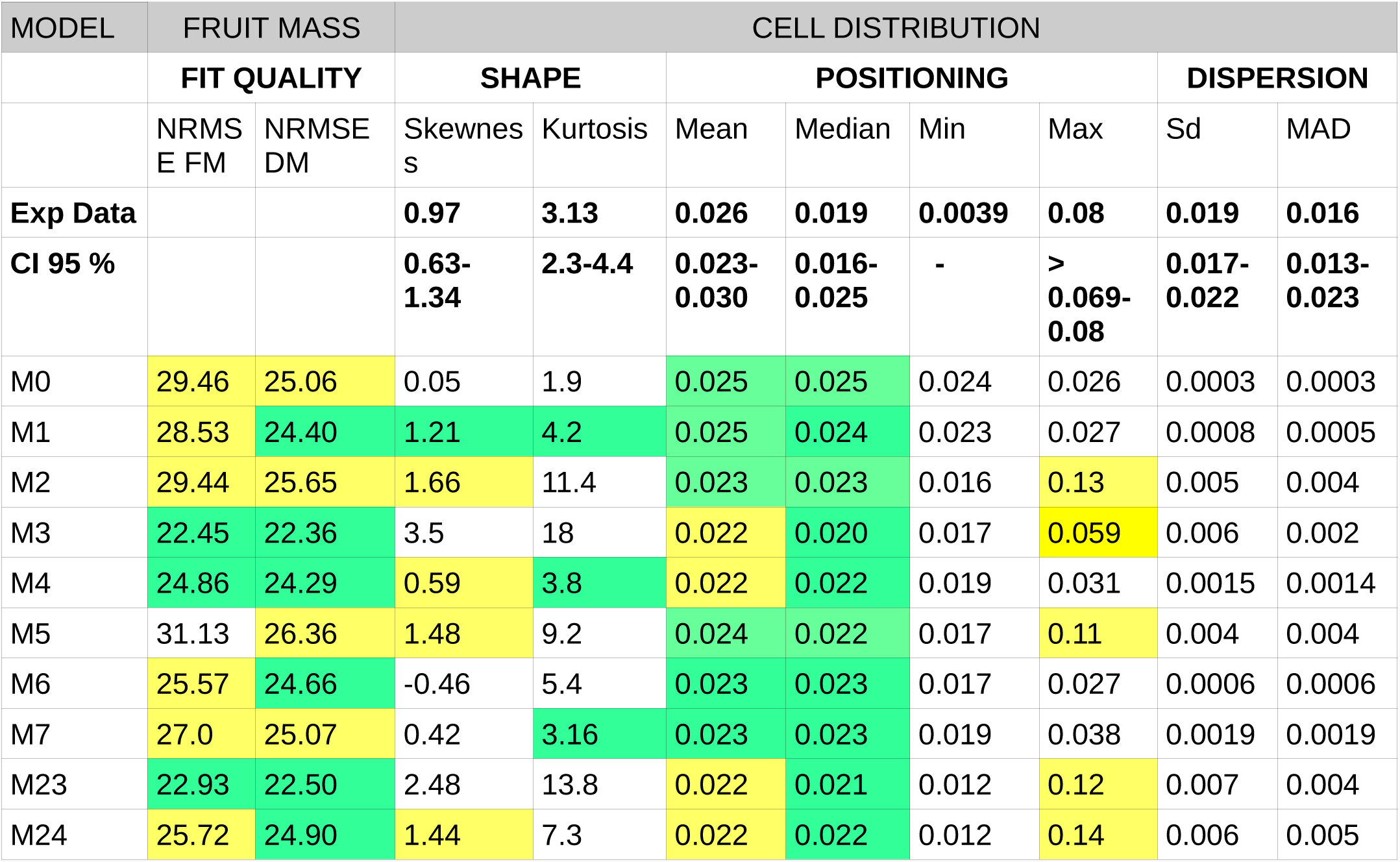
Statistical descriptors for the measured and predicted cell area distribution for Cervil genotype. The NRMSE scores for predicted pericarp dry and fresh masses corresponding to the selected solution are reported under the columns “Fit Quality”. For an easier interpretation, green and yellow boxes indicate, respectively, a satisfactory (i.e. NRMSE < 25 and a moment values within the confidence interval of experimental measurement) and a partial agreement to data (NRMSE between 25 and 30, moments near the limit of the confidence interval).

**Table 3:**
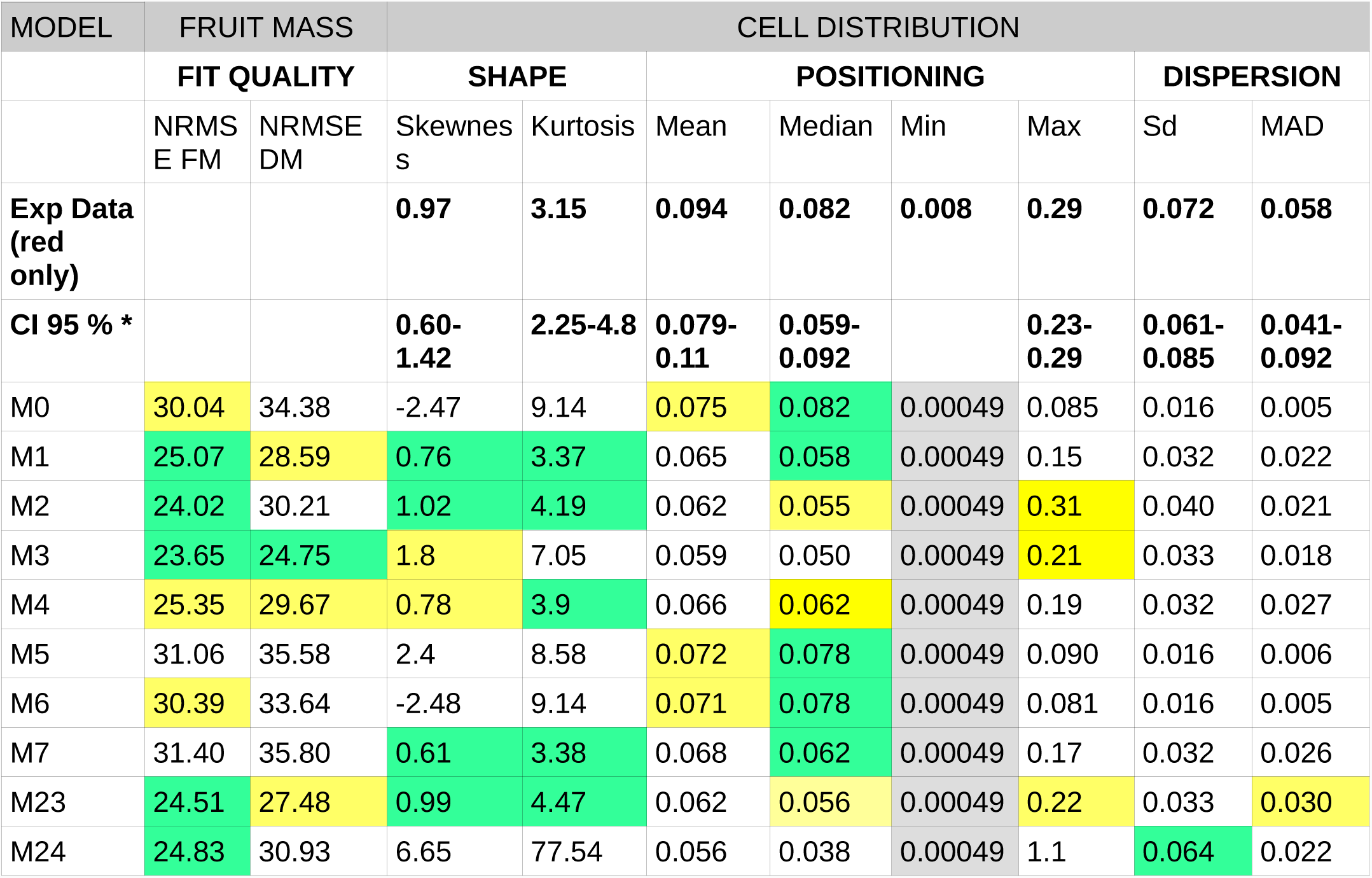
Statistical descriptors for the measured and predicted cell area distribution for Levovil genotype. The NRMSE scores for predicted pericap dry and fresh masses corresponding to the selected solution are reported under the columns “Fit Quality”. For an easier interpretation, green and yellow boxes indicate, respectively, a satisfactory (i.e. NRMSE < 25 and a moment values within the confidence interval of experimental measurement) and a partial agreement to data (NRMSE between 25 and 30, moments near the limit of the confidence interval).

| MODEL | FRUIT MASS |  | CELL DISTRIBUTION |  |  |  |  |  |  |  |
| --- | --- | --- | --- | --- | --- | --- | --- | --- | --- | --- |
|  | FIT QUALITY |  | SHAPE |  | POSITIONING |  |  |  | DISPERSION |  |
|  | NRMSE FM | NRMSE DM | Skewness | Kurtosis | Mean | Median | Min | Max | Sd | MAD |
| **Exp Data (red only)** |  |  | **0.97** | **3.15** | **0.094** | **0.082** | **0.008** | **0.29** | **0.072** | **0.058** |
| **CI 95 % \*** |  |  | **0.60-1.42** | **2.25-4.8** | **0.079-0.11** | **0.059-0.092** |  | **0.23-0.29** | **0.061-0.085** | **0.041-0.092** |
| M0 | 30.04 | 34.38 | -2.47 | 9.14 | 0.075 | 0.082 | 0.00049 | 0.085 | 0.016 | 0.005 |
| M1 | 25.07 | 28.59 | 0.76 | 3.37 | 0.065 | 0.058 | 0.00049 | 0.15 | 0.032 | 0.022 |
| M2 | 24.02 | 30.21 | 1.02 | 4.19 | 0.062 | 0.055 | 0.00049 | 0.31 | 0.040 | 0.021 |
| M3 | 23.65 | 24.75 | 1.8 | 7.05 | 0.059 | 0.050 | 0.00049 | 0.21 | 0.033 | 0.018 |
| M4 | 25.35 | 29.67 | 0.78 | 3.9 | 0.066 | 0.062 | 0.00049 | 0.19 | 0.032 | 0.027 |
| M5 | 31.06 | 35.58 | 2.4 | 8.58 | 0.072 | 0.078 | 0.00049 | 0.090 | 0.016 | 0.006 |
| M6 | 30.39 | 33.64 | -2.48 | 9.14 | 0.071 | 0.078 | 0.00049 | 0.081 | 0.016 | 0.005 |
| M7 | 31.40 | 35.80 | 0.61 | 3.38 | 0.068 | 0.062 | 0.00049 | 0.17 | 0.032 | 0.026 |
| M23 | 24.51 | 27.48 | 0.99 | 4.47 | 0.062 | 0.056 | 0.00049 | 0.22 | 0.033 | 0.030 |
| M24 | 24.83 | 30.93 | 6.65 | 77.54 | 0.056 | 0.038 | 0.00049 | 1.1 | 0.064 | 0.022 |

### A simple cell-autonomous control scheme leads to unrealistic cell size distribution

As a benchmark model, the case of a simple cell-autonomous control, without ploidy-dependent effect, was first considered (version M0 of the model). Accordingly, two cells with the same age, even if initiated at different fruit developmental stages, behave identically in what concerns carbon metabolism, transport and wall mechanical properties. In this scheme, therefore, cell size variations derive exclusively from the dynamics of cell division, that cause a shift in the initiation date for different cohort of cells. When applied to our genotypes, the cell-autonomous model was able to reproduce the observed pericarp mass dynamics but the corresponding cell size distribution was extremely narrow (see Table 2 and 3).

Including an organ-wide mechanism that controls cell size (model M1) introduces a source of variance among cells. In this case, two cells of the same age which were initiated at different fruit stages do *not* behave identically, resulting in different expansion capabilities and growth patterns (Figure 3, Baldazzi et al. 2013). As a result, standard deviation doubled and skewness increased towards small positive values, indicating a slightly right-tailed cell size distribution, both for both cherry and large-fruited tomatoes (Table 2 and 3). However, the maximum cell size predicted by the model remained much smaller then expected, suggesting that a mechanism controlling cell expansion is lacking in the model.

**Figure 3:**
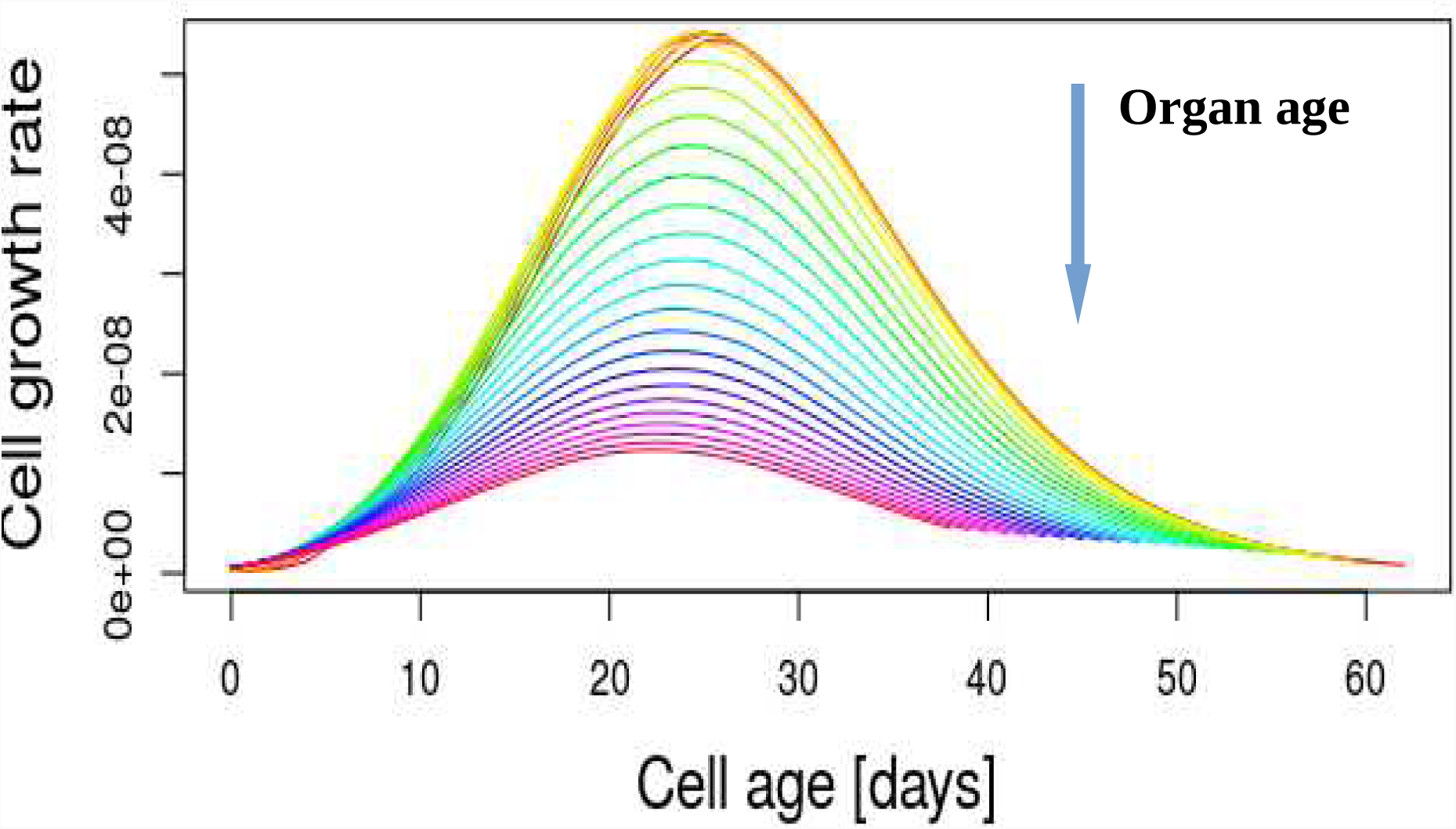
Effect of the organ-wide control on the cell growth rate. Cells that enter the expansion phase late during organ development (large organ age) have a lower growth rate due to the progressive reduction of the symplastic carbon transport. Simulations have been obtained with model M1, Levovil genotype.

### Endoreduplication and cell growth: possible action and genotypic effect

The suggestion that nuclear ploidy level may be important for cell size control has been often reported in litterature. However, the molecular mechanism by which ploidy could modulate cell expansion capabilities remain elusive. In this work, three cell properties have been selected as possible targets of ploidy-dependent modulation (see M&M section): *a)* the maximum carbon uptake rate *b)* carbon allocation between soluble and non-soluble compounds *c)* cell wall plasticity. The three hypotheses were tested on both genotypes, in combination or not with an organ-wide control.

A principal component analysis was performed on 8 statistical descriptors of cell size distribution in order to compare predictions of the different models (see M&M and Tables 2 and 3). For both genotypes, the first two principal components explained approximately 90% of observed variance (Figure 4 and 5). Separation was mainly performed by the first principal component on the basis of the width of the distribution (sd, mad and maximal cell size), on one side, and its mean and median values, on the other direction.

**Figure 4:**
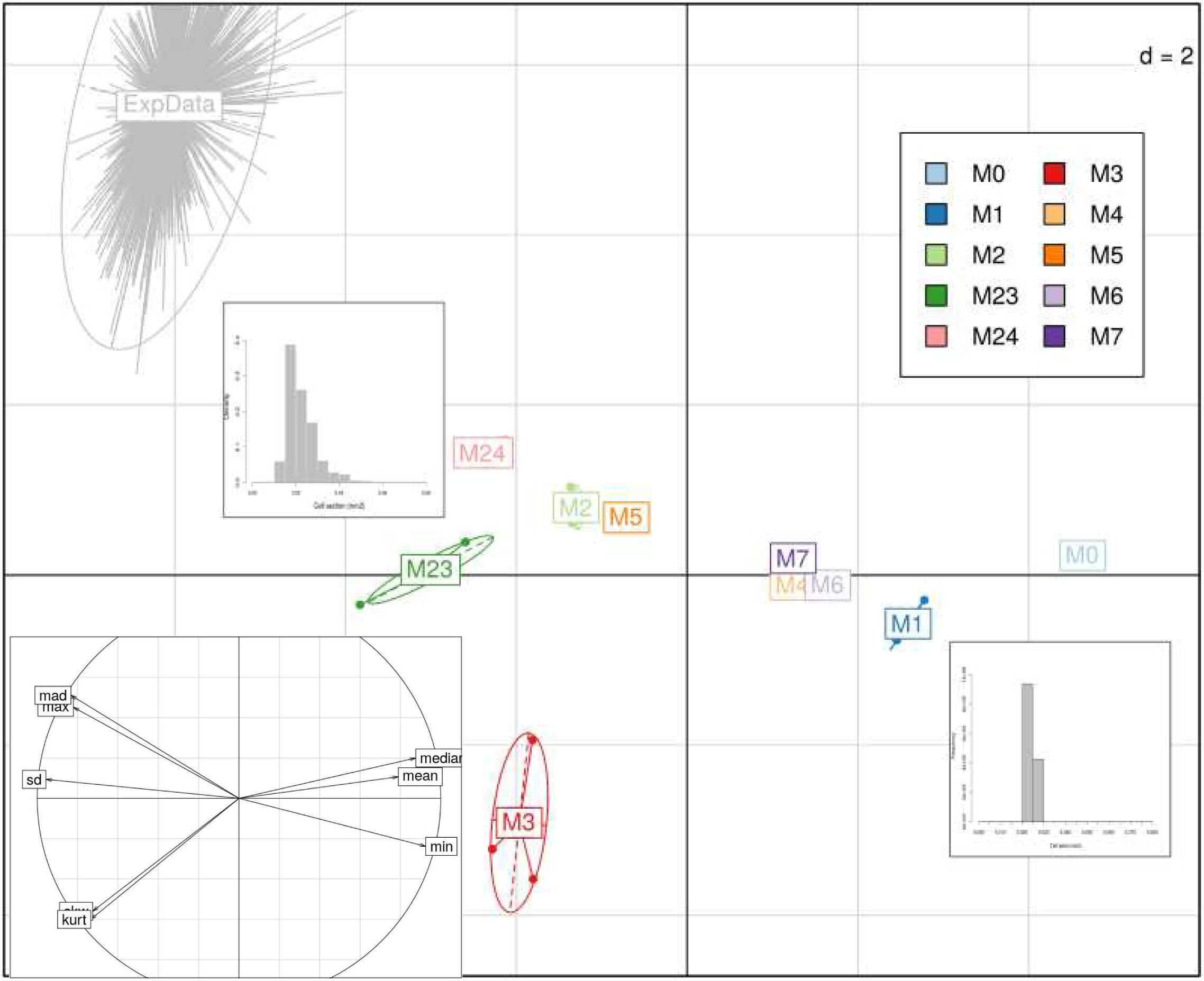
Principal component analysis (PCA) cell size distributions obtained for the different estimations of models M0-M24 on Cervil genotype. *Main plot:* Projection of individual distributions on the PC1-PC2 plane (respectively 72% and 16% of variance explained); *d* is the grid unit. Bootstrap results on measured cell size data are projected as a supplementary observation. Model variants are tagged with different colors. Typical cell size distribution shapes are sketched for the main subgroups. *Inset:* Correlation of the variables with the first two principal components.

**Figure 5:**
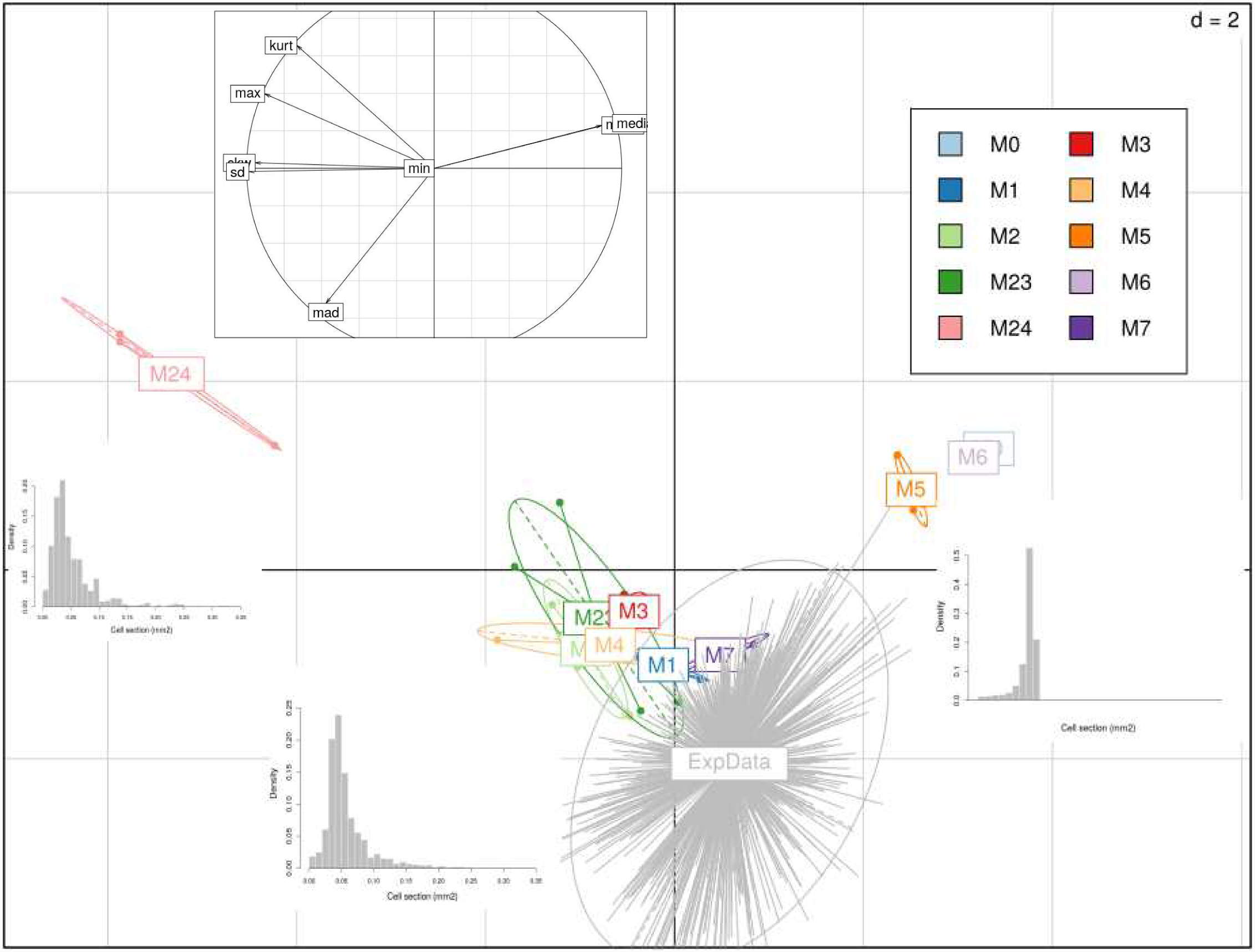
Principal component analysis (PCA) cell size distributions obtained for the different estimations of models M0-M24 on Levovil genotype. *Main plot:* Projection of individual distributions on the PC1-PC2 plane (respectively 75% and 17% of variance explained); *d* is the grid unit. Bootstrap results on measured cell size data are projected as a supplementary observation. Model variants are tagged with different colors. Typical cell size distribution shapes are sketched for the main subgroups. *Inset:* Correlation of the variables with the first two principal components.

As already mentioned, models with simple cell-autonomous control were characterized by narrow distributions but centered around a larger mean (and median) value. The addition of an organ or ploidy effect on cell expansion resulted in an increase in cell size variance and skewness, shifting the distribution towards the observed right-tailed shape (Table 2 and 3). Models combining an organ-wide and a ploidy-dependent control were closer to experimental data, although they could not fully approach the observed distribution in the case of the cherry tomato genotype.

When analysed in details, results show that the relative importance of the organ-wide and the ploidy-dependent control of cell expansion was genotype-dependent. In the case of Levovil, organ-wide control turned out to be the major regulatory mode. With the exception of M7, models without organ-control (models M0, M5,M6) completely failed to reproduce the observations, resulting in a very narrow and left-tailed cell size distribution. Organ-wide coordination of cell expansion appeared to be the main responsible for positive skewness of cell size distribution whereas the addition of an endoreduplication-mediated modulation of cell expansion capabilities, alone, resulted only in a marginal improvement of model’s performances.

The relative roles of ploidy-dependent and organ-wide control of cell growth appeared more balanced in cherry tomatoes. Both models including an organ-wide (M1) *or* a ploidy-mediated control of cell expansion (models M5 to M7) resulted in very narrow and quite symmetric distribution. The concomitant action of both control mechanisms was needed in order to get the expected right-tailed distribution and realistic cell size variations (model M2-M4, M23 and M24). The two mechanisms thus seem to act in synergy to increase cell expansion and final cell size.

### A direct influence of endoreduplication on cell expansion is needed to get correlation between cell size and ploidy

A strong correlation has been often reported between cell size and ploidy level. This may be partly innate to the temporal evolution of endoreduplication, as cells with a high ploidy necessary had more time to growth, without being halved by cell division. On this basis, some authors have claimed that the observed correlation between size and ploidy is just a matter of time and there’s no need of any direct effect of endoreduplication on cell growth to explain the data (Roeder *et al*., 2010; Robinson *et al*., 2018).

We checked this intuition with the help of our modelling framework. A linear regression analysis between cell size and DNA content was performed for all tested models, at fruit maturity. Results are reported in table S5.

For both genotypes, no correlation was found for models M0 and M1 (Figures 6 and 7, left panel) due to the asynchrony in cell division and endoreduplication patterns. Indeed, cells can attain a same ploidy level following different temporal sequences of expansion and endocycle events, leading to a large variability of possible cell sizes and ages within a same ploidy class. As long as ploidy level and growth rate are independent (as in models M0 and M1), the specific endoreduplication pattern has no consequence on the final cell size: variations in cell sizes simply reflect variations in cell ages and no correlation is found, on average, between the ploidy and size.

**Figure 6:**
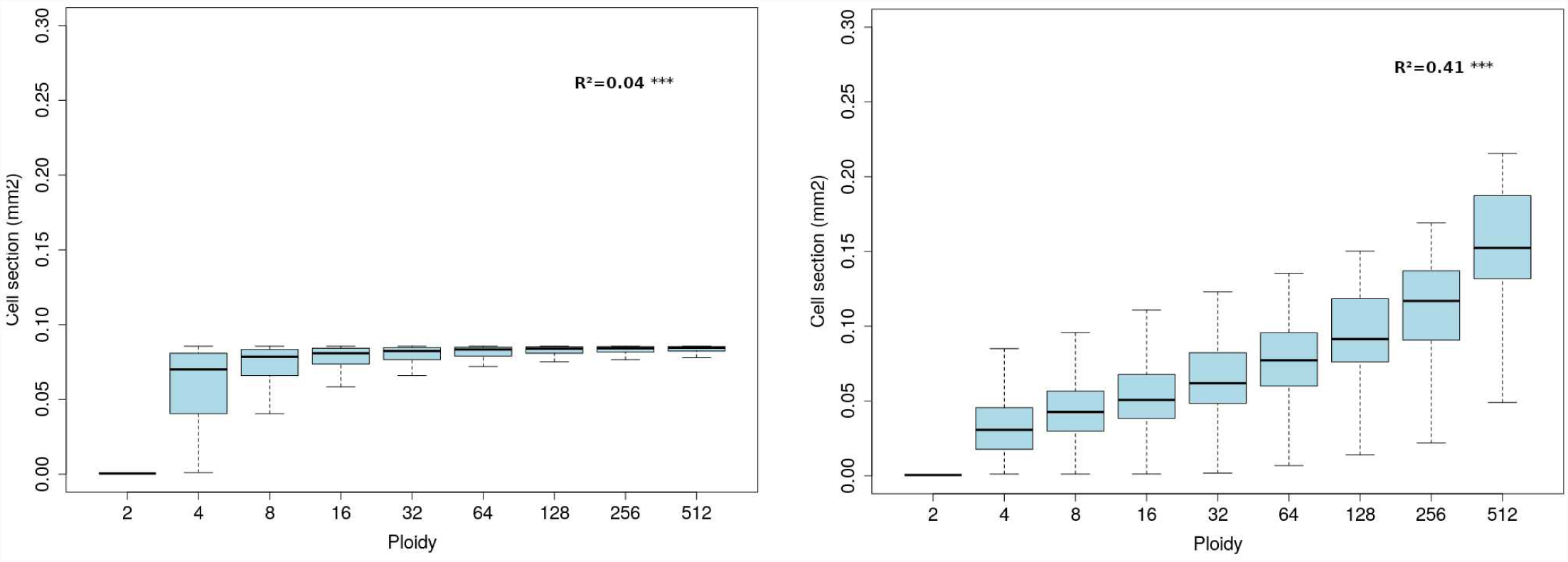
Simulated relation between ploidy and cell size at fruit maturity for the Levovil genotype. *Left:* model M0. *Right:* model M23. The adjusted R^2^ corresponding to a linear regression model is reported.

**Figure 7:**
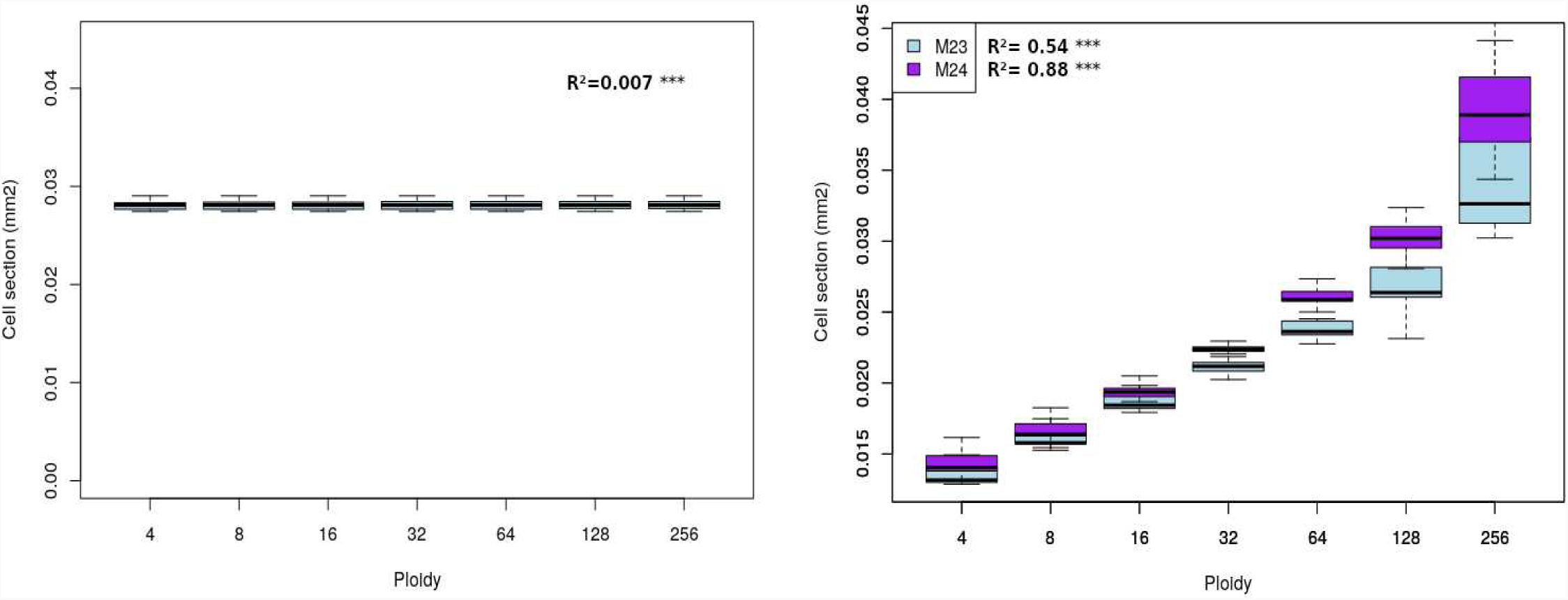
Simulated relation between ploidy and cell size at fruit maturity for the Cervil genotype. *Left*: model M0. *Right*: models M23 and M24. The adjusted R^2^ corresponding to a linear regression model is reported.

A direct effect of ploidy on cell expansion rate was needed in order to get a non-zero correlation between ploidy and size in our model. In particular, models in which the ploidy level affected the carbon import rate (M2, M5, M23, M24) led to a significant positive correlation between ploidy and size (p-values <0.001), for both genotypes (see Table S5, Figure 6 and 7, right panel). In these models, the observed increase in cell size with increasing ploidy level was directly linked to enhanced cell expansion capabilities, and significant correlation was found between ploidy and maximal cell growth rate (Table S6).

Interestingly, the heterogeneity in cell sizes increased with increasing ploidy (Levene test on size variance, p-value <0.001), in agreement to what observed in another tomato variety (Bourdon et al 2011). Cell size variations were larger for Levovil genotype than for the cherry tomato variety due to its extended division phase, that increased variability in the timing of exit from the mitotic phase (Figure 6, right panel).

## DISCUSSION

The present paper describes an improved version of an integrated cell division-expansion model that explicitly accounts for DNA endoreduplication, an important mechanism in tomato fruit development. The model is used to investigate the interaction among cell division, endoreduplication and expansion processes, in the framework of the neo-cellular theory (Beemster et al., 2003).

To this aim, 10 model variants including or not a ploidy-dependent and an organ-wide control of cell development have been tested and compared to data from two contrasting tomato genotypes. Specific cellular processes have been hypothesized as possible targets of both modes of control, based on literature information. It is important to stress that the molecular basis of the supposed regulations are not described in the model and could involve many molecular players, including hormones, mechanical signals etc. Moreover, the existence of other targets for organ-wide or ploidy-dependent regulations cannot be excluded, as well as the contribution of other mechanisms to the control of cell growth. The objective of the paper, indeed, was not to identify the exact mechanism of interaction between endoreduplication and expansion, but rather to test *if* a direct influence of ploidy onto cell expansion, in combination or not with an organ-wide control, was likely to be involved in the control of fruit growth.

Model simulations showed that a pure cell-autonomous control was unable to reproduce the observed cell size distribution, resulting in a very narrow distribution, well different from the expected skewed, right-tailed shape. In agreement with the neo-cellular theory, the model supports the need for an organ-wide control of cell growth as a key mechanism to increase cell size variance and points to a direct effect of cell ploidy on cell expansion potential.

### Measurement of cell size distribution: a promising approach to understand the control of fruit growth

Our work is based on the analysis of cell size distribution as a footprint of different control schemes. When looking at our results, indeed, the NRMSE with respect to pericarp fresh and dry mass data was always between 20% and 30% indicating a satisfactory agreement with data, independently from the model version and the tomato genotype. This highlights the fact that the dynamics of fruit growth alone is not enough to discriminate between several biologically-plausible models. In this sense, cell size distribution represents a much more informative dataset as it uniquely results from the specific cell division and expansion patterns of the organ (Halter et al., 2009).

The assessment of cell sizes in an organ is not an easy task though and the employed measurement technique may have important consequences on the resulting cell size distribution (Legland et al., 2012). Indeed, mechanical constraints acting on real tissues as well as vascularisation can largely modify cell shape, resulting in elongated or multi-lobed cells (Ivakov and Persson, 2013). Thus, if the orientation of 2D slices can potentially affect the resulting cell area estimation, possible differences between *in-vivo* tissues and dissociated cells should also be systematically checked (McAtee et al. 2009). The size of the dataset is also important to correctly characterized the expected distribution shape. Indeed, outliers can significantly affect the estimation of high-order moments, especially in heavy-tailed distribution like the one usually observed in plant organs. The above reasons explain why we decided to focus on a qualitative comparison of simulated and experimental cell size distribution rather than on a perfect fit. Moreover, uncertainty in our dataset has been accounted for via the the estimation of confidence intervals for the experimental distribution moments.

In perspective, the use of mutant or modified strains (Musseau et al., 2017) in combination with recent advancements in microscopy and tomography could permit the acquisition of more reliable datasets, opening the way to a in-depth investigation of cell size distribution in relation to fruit tissues and to the underlying molecular processes (Mebatsion et al., 2009; Wuyts et al., 2010).

### The relative importance of organ-wide and ploidy-dependent controls may be genotype-dependent

According to the model, organ-wide control was responsible for cell-to-cell variations but a ploidy-mediated effect on cell expansion was needed in order to obtain a significant correlation between size and ploidy as observed in experimental data of fruit pericarp (Bourdon et al., 2011). However, the relative importance of the two modes of control may be genotype-dependent.

For the large-fruited variety, organ-wide control was the dominant mechanism. This is probably due to its long division phase that causes the appearance of new expanding cells late in the fruit development, once plasmodesmata closure is already completed. Independently of the targeted process, the addition of a ploidy-dependent effect instead, did not significantly modify the predicted cell size distribution. The model supports the idea that cell ploidy may fix a maximum potential growth rate. Given the large fruit mass in Levovil genotype, it is possible that such a potential may not be completely reached in our experimental conditions, due to limited plant resources.

In the case of the cherry tomato variety, the effect of a ploidy-dependent mechanism was more pronounced, especially when affecting cell carbon metabolism. Models combining both an organ and a ploidy-dependent control performed better then the others although they failed to fully account for the experimental cell size distribution. A few reasons may explain this discrepancy. First, due to a lack of data, the endoreduplication dynamics has been calibrated on the 2004 experiment whereas the division and the expansion modules have been estimated on the 2007 data, for which the cell size distribution was available. Little is known about the possible dependence of endoreduplication on environmental variables (Engelen-Eigles *et al*., 2000; Setter and Flannigan, 2001; Cookson *et al*., 2006). In tomato fruit, changes in ploidy levels were mainly linked to changes in the duration of the mitotic phase (Bertin, 2005) but a direct effect of environmental fluctuations on endoreduplication-related parameters cannot be excluded. This may be particularly true for the cherry tomato genotype, for which the dynamics of cell division differed significantly between the two years (see Supplemental data, section S3). One can therefore expect that the progression of endocycle may have been different too, with possible consequences on the shape of the resulting cell size distribution. Preliminary simulations showed that an acceleration of the endocycle or an increase of the proportion of cells that enter a new round of endoreduplication can spread the resulting distribution towards large cell sizes, increasing the overall variance in models including a ploidy-dependent effect.

In addition to cell-cycle-related mechanisms, environment and cultural practices can also affect resources availability at the cell scale. In many fruit species including tomato, a negative correlation between average cell size and cell number has been observed, suggesting the existence of a competition for resources (Prudent et al., 2013). This kind of mechanism may widen the range of attainable cell sizes, increasing size variations between first and late-initiated cells. The importance of such an effect may vary with genotype and environmental conditions (Bertin, 2005; Quilot and Génard, 2008).

### Stochasticity in cellular processes may be important to explain cell size variance in fruit

Our model is an example of population model: the fruit is described as a collection of cell groups, each having specific characteristics in terms of number, mass, age and ploidy level, that dynamically evolve during time. Although asynchrony in the emergence of cell groups allowed to capture a considerable part of cell-to-cell heterogeneity, the intrinsic stochasticity of cellular processes (Meyer and Roeder, 2014; Robinson et al., 2011b; Smet and Beeckman, 2011) is not accounted for. Variations in the threshold size for division (often associated to a change in the cell cycle duration) as well as asymmetric cell divisions are considered as important determinant of the final cell size (Dupuy et al., 2010; Osella et al., 2014; Roeder et al., 2010; Stukalin et al., 2013). They may contribute to significantly spread the size distribution of both proliferating and expanding cell groups, from the early stages. Moreover, the degree of additional dispersion introduced by cell expansion is likely to depend on the specificity of the underlying mechanisms, with possible interactions with ploidy-dependent and organ-wide controls.

In perspective, the addition of stochastic effects could help to fill the missing variance for both Cervil and Levovil genotypes. To this aim, a novel modelling scheme is needed in which the average cell mass of a group is replaced by a distribution function of cell sizes, whose parameters can evolve with time under the effect of cell expansion processes.

### Correlation between size and ploidy: a clue for a direct influence of endoreduplication on cell expansion in tomato fruit?

Our results showed that asynchrony in cell division and endoreduplication events prevented the emergence of a correlation between size and ploidy based only on time proceeding. According to the present model, a direct effect of nuclear ploidy on the attainable cell growth rate is needed in order to obtain the observed correlation in tomato fruit. This is in line with literature data pointing to ploidy level setting the maximum cell growth rate that can be attained or not, depending on internal (hormones) and external (environmental) factors (Breuer et al., 2010; Chevalier et al., 2011; De Veylder et al., 2011). Of course, this result may be less striking in systems where the progression of endocycles is more sequential. This may be the case of *Arabidopsis thaliana* sepals, where data are consistent with a model in which expanding cells undergo a new round of endoreduplication at each time step (Roeder et al., 2010). In this latter system, variability among cell size arises from asymmetry in cell division and variations in the exit time from mitotic cycle, whereas no differences in the growth rate is observed among cells with different ploidy level (Tauriello *et al*., 2015; Robinson *et al*., 2018).

Overall, these results confirm that the relationship between endoreduplication and expansion may not be universal but can differ depending on the considered organ. Moreover, attention should also be payed to cell identity. Indeed, quantitative differences in the strength of the ploidy-dependent effect have been demonstrated between pavement and mesophyll cells in *Arabidopsis* leaves (Katagiri et al., 2016; Kawade and Tsukaya, 2017), whereas cell-layer specific developmental patterns have been observed in tomato fruit (Renaudin et al. 2017). In the model developed here, the spatial distribution of expanding-endoreduplicating cells is not accounted for. In perspective, a model including distinct cell-layer populations, with specific expansion programs, may help to refine the relation between ploidy-size.

At term, improvements in the ability of computation models to integrate the multiple facets of organ development in a mechanistic way can help to evaluate and quantify the contribution of the different processes to the control of cell growth.

## Acknowledgments

The authors warmly thank B. Brunel for help with cell measurements. The authors are grateful to Inria Sophia Antipolis – Méditerranée “NEF” computation cluster for providing resources and support. This work was partially funded by the Agence Nationale de la Recherche, Project “Frimouss” (grant no. ANR–15– CE20–0009) and by the Agropolis Foundation under the reference ID 1403-032 through the « Investissements d’avenir » programme (Labex Agro:ANR-10-LABX-0001-01).

