## Supplemental File for "Organ-wide and ploidy-dependent regulations both contribute to cell size determination: evidence from a computational model of tomato fruit"

### S1 Cell Distribution fit

An approximate Kolmogorov-Smirnov test has been performed using R library ‘fitdistriplus’ (gofstat function) in order to establish the shape of our cell-size distribution. Results showed that experimental data are compatible both to gamma and weibull distributions, at the significance level 0.05, for both genotypes (Fig S1 and S2). Parameters of the corresponding gamma and weibull fit are reported in table S1.

Note however that approximate KS test assumes the distribution parameters known and thus tends to be quite conservative with respect to other statistical tests. For our datasets, both the Anderson-Darling and Kramer-von Mises tests showed not significant at a p-value of 0.05.

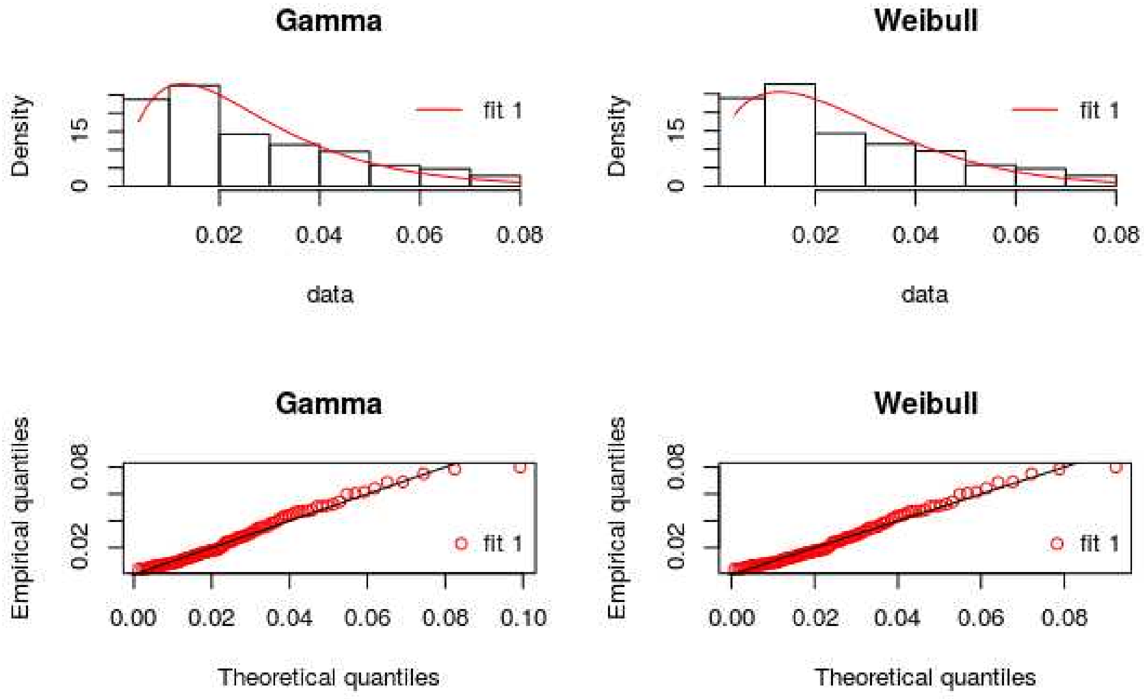

**Figure S1:** Fit of the cell size distribution for Cervil genotype. *Left:* Fit using a Gamma distribution function. *Right:* Fit using a Weibull distribution function. Theoretical and experimental density functions are compared in the top part of the graph, whereas the corresponding quantiles are reported in the bottom panels.

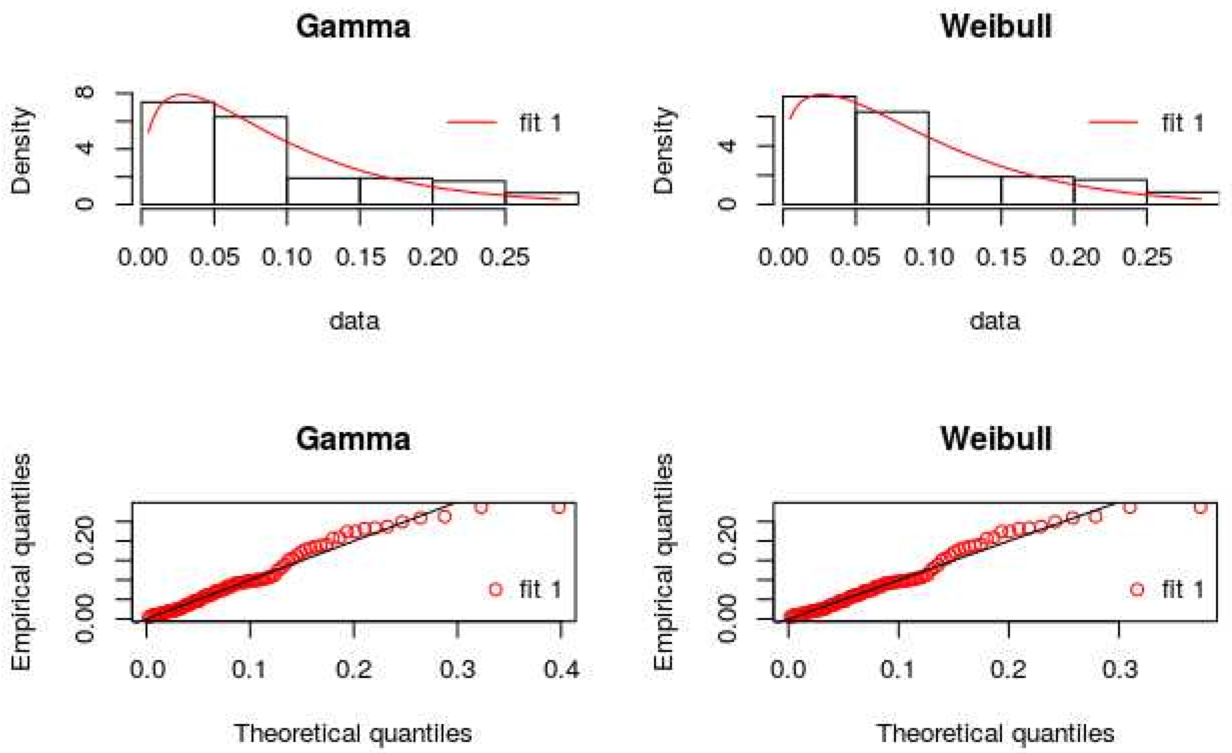


**Figure S2:** Fit of the cell size distribution for Levovil genotype. *Left:* Fit using a Gamma distribution function. *Right:* Fit using a Weibull distribution function. Theoretical and experimental density functions are compared in the top part of the graph, whereas the corresponding quantiles are reported in the bottom panels.

| | Gamma Distribution | | Weibull Distribution | |
|---|---|---|---|---|
| | shape | rate | shape | scale |
| **Cervil** | 1.94 +- 0.25 | 74.25+- 10.83 | 1.45 +-0.11 | 0.029 +- 0.00021 |
| **Levovil** | 1.44+-0.19 | 15.69+-2.46 | 1.25 +- 0.1 | 0.099 +- 0.0086 |

**Table S1**: Distribution parameters of the Gamma and Weibull distribution fit for the two genotypes.

### S2: Implementation of the integrated model

The integrated division-endoreduplication-expansion model has been implemented in Java as an object-oriented model.

Two cell classes are defined: the proliferating cells and the expanding (and endoreduplicating) cells. At time zero all cells are initialized as proliferating cells.

Proliferating cells are characterized by their number, whereas they are assumed to have a constant mass *m0* and ploidy level 2C.

There is a  single group of proliferating cells. Growth of proliferating cells within two successive division events is not modeled explicitly.  At each mitotic cycle *tau*, a fraction of proliferating cells proceed through division whereas the remaining ones enter the "expanding cells " class. Cell divisions are implicitly assumed to be symmetric, each (proliferating) daughter cell having exactly the same mass *m0* as their mother.

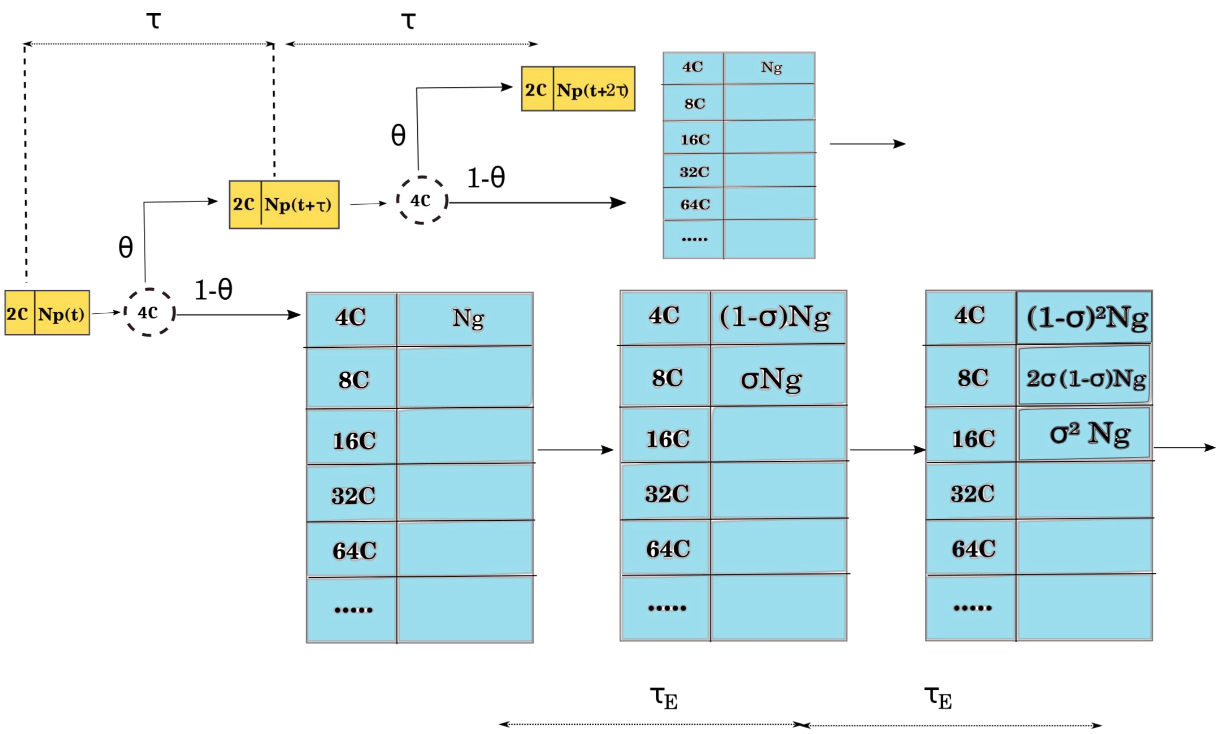


**Figure S2:** Schematic representation of the structure of an expanding cell group as an array of  sub-classes  corresponding to possible ploidy levels.  At initialization,  all expanding cells  are put into the 4C ploidy. Every endocycle  a fraction  $\sigma$  of the cells  increase its ploidy level *p* by a factor 2, moving to the following ploidy class  whereas the remaining ones keep their initial ploidy *p*.

### *S2.1  Division-endoreduplication module*

The division-endoreduplication module governs the evolution of the number of cells in each classes and ploidy level.  With respect to the orginal model (Bertin et al., 2007), the version implemented here is slightly simplified as it does not consider the presence of inactive cells that neither divide nor endoreduplicate.  Accordingly, in each mitotic cycle

$\tau$ , the proportion of cells proceeding through division $N_{p(}$(t) depends only on the the proliferating capacity $\theta(t)$ , which progressively declines during fruit development:

$$N_p(t)=2*N_p(t-1)*\theta(t)$$

The model assumes that cells that do not divide switch to expansion *and* to repeated syntheses of DNA without cell division, resulting in cell endoreduplication. Thus, expanding cells are characterized by their number, mass and ploidy level. Any expanding cell group is structured as an array of sub-classes corresponding to possible ploidy levels from 4C to 512 C (Levovil genotype) or 4C to 256C (Cervil genotype). All cells of a same group have the same age, but each subclass has a proper cell number and cell mass (when a ploidy effect is included into the model).

Unlike proliferating cells which are pooled in a single group, a new group of $N_g$(t,p) expanding cells is created, at each mitotic event, together with its sub-classes of possible ploidy levels *p*. At initialization, all expanding cells of the group are put into the 4C ploidy class, with mass 2*m0:*

$$N_g(t,4)=N_p(t-1)(1-\theta(t))$$

The remaining ploidy class are empty.

A single constant parameter $\sigma$ describes the proportion of cells that moves from one to the next class of DNA content after each lapse of time, $\tau_E$ , considered to be the minimum time required for an endocycle. Every time $\tau_E$ , in each group of expanding cells, a fraction $\sigma$ of the cells increase its ploidy level *p* by a factor 2, moving to the following ploidy sub-class:

$$N_g(t,2p)=\sigma N_g(t-1,p)$$

The remaining cells of the group keep the initial ploidy level *p*

$$N_g(t,p)=(1-\sigma)N_g(t-1,p)$$

Once the number of cells in each ploidy class updated, their mass is recalculated as a weighted average of its components. The mass of expanding cells increases at each time step according to the expansion module and the available resources from the plant, as explained in the next section.

*S2.2 Expansion module*

Following (Baldazzi et al., 2013; Fishman and Génard, 1998; Liu et al., 2007) cell is described as a compartment that takes up water and sugar through a composite membrane, separating it from the xylem and phloem, and loses water and dry matter through the processes of transpiration and respiration.

The cell state at any time is described by two state variables, the mass of water (w (g)) and the dry weight (s (g)). The model is driven at an hourly time step by four input variables. Two of these are properties of the external environment: humidity (H) and temperature (T ($^oC$)) of the air. The other two are properties of the vasculature: the water potential of the vasculature ($\psi_x$ (bar)) and the concentration of sugars in the phloem ($C_p$ (g $g^{-1}$)). It is assumed that the water potential of the phloem is the same as that of the xylem, as the separating membrane is highly permeable to water, so their hydrostatic pressures differ only due to differences in solute potentials.

In brief, the model of Fishman and Génard can be described as follows. The rates of change of fruit water (w) and dry matter (s) at any time (t) are given by

$$\frac{dw}{dt} = U_x + U_p + r_w R_c - T_c$$

$$\frac{ds}{dt} = U_s - R_c$$

where $U_x$ and $U_p$ are the amounts of water taken up per unit time from xylem and phloem respectively, $U_s$ is the dry matter uptake rate, and $T_c$ and $R_c$ are **cell** transpiration and respiration rates respectively. Note that following Brussières (1993) we have assumed that a fraction $r_w$ =0.6 of respired dry matter is converted to water. In the absence of any geometrical description of cell position within the fruit, cell transpiration rate is computed as a fraction of the total fruit transpiration $T_f$ as

$$T_c = \frac{w}{W_f} T_f$$

where $W_f$ is the total fruit water. As in the original models, fruit transpiration is driven by the difference between the humidity of air spaces within the fruit ($H_f$ = 0.996 as in Fishman and Génard, 1998) and the humidity of the air at the fruit surface Af:

$$T_f = A_f \rho \alpha(T)(H_f - H)$$

where ρ is the permeation coefficient of the fruit surface to water vapour (cm h -1 ), and $\alpha(T)$ is a function of the external temperature (see Fishman & Génard for more details). Following (Baldazzi et al., 2013), the permeation coefficient ρ decreases with fruit age ($t_f$) due to an increased deposition of wax layer at the organ surface (Lee, 1990), as

$$\rho=\rho_1+\rho_0 e^{-(k_\rho(t_f-t0))}$$ .

We denote the osmotic and hydrostatic pressures in the phloem vasculature by $\pi_p$ and $P_p$ ($=\pi_p+\psi_x$) respectively, and in the xylem by $\pi_x$ and $P_x$. The equations used to describe the mass flow through the composite membrane are the same as those used by Fishman and Génard (1998),

$$U_x=A_x L_x[P_x-P_c-\sigma_x(\pi_x-\pi_c)]$$

$$U_p=A_p L_p[P_p-P_c-\sigma_p(\pi_p-\pi_c)]$$

where $P_c$ and $\pi_c$ are the cell turgor and osmotic pressure, respectively. The osmotic pressure in the xylem vasculature is set to zero, and as the plasma membrane is largely impermeable to sugars, a reflection coefficient $\sigma_x$ is assumed 1.

Following (Liu et al., 2007) the effective reflection coefficient $\sigma_p$ of the membrane separating the phloem vessel (plasmodesmata) from cell is assumed to be dependent on time as

$$\sigma_p=1-\exp(-\tau_s t^2)$$

The reflection coefficient thus passes from a permeable membrane ( $\sigma_p=0$ ) to a fully impermeable barrier as the time t increases, thus controlling the switch between symplastic and apoplastic carbon transport (see eqn for $U_s$). For model versions assuming an organ-level control, *t* corresponds to the age of the fruit, $t_f$..

Uptake of sugars (and hence dry matter) from the phloem into the fruit ($U_s$) has three components:

$$U_s=U_a+\sigma_p C_s U_p+A_p p_s(C_p-C_c)$$

The first term is the active uptake, the second term is uptake due to the mass flow above (symplasmic transport), and the third term is diffusive flow given a total permeability of the membrane . $C_p$ and $C_c$ are the concentrations (proportions by weight) of sucrose in the phloem vasculature and cell respectively,  and $C_s$ is the average of these two. It is assumed that a proportion ssrat of the dry matter s is in soluble form, i.e.

$$C_s = \frac{ssrat * s}{w + ssrat * s}$$

where ssrat is a function of the age of the cell $t_c$ (Baldazzi et al., 2013)

$$ssrat = b_{ssrat}\left(1 - e^{-a_{ssrat} * t_c}\right) + ssrat_0$$

Active uptake Ua is described by Michaelis-Menten kinetics as

$$U_a = \frac{\nu_m s\, C_p}{\left(K_M + Cp\right)} \frac{1}{\left(1 + e^{\tau_a\left(t_c - t_{star}\right)}\right)}$$

where $\nu_m$ is the maximum uptake rate per cell dry weight (g $h^{-1}$ ) and $K_M$ is the Michaelis constant. The second factor  includes the effect of  a non-competitive inhibition late in cell development, as in the original paper by Fishman & Génard (1998).

Turgor $P_c$ is calculated by equating two expressions for the rate of change of the volume of the cell. Cell volume (V) can be written simply as

$$\frac{dV}{dt} = \frac{1}{D_s}\frac{ds}{dt} + \frac{1}{D_w}\frac{dw}{dt}$$

where $D_w$ (=1) and $D_s$ (=1.6) are the densities of water and carbohydrate respectively.  The second expression used is Lockhart's equation (Lockhart, 1965)

$$\frac{dV}{dt} = V\,\phi\left(P_c - Y\right)$$

where $\phi$ is the cell's plasticity and Y the turgor threshold. Following (Liu et al., 2007) membrane plasticity is assumed to decrease during cell development as:

$$\phi = \phi_{min} + \frac{\left(\phi_{max} - \phi_{min}\right)}{1 + e^{k\left(t_c - t_0\right)}}$$

By equating the two expression for the rate of volume change, an algebraic expression for $P_c$ can be obtained.

### S3: Model calibration and estimated parameters values

#### *S3.1 Calibration of the division-endoreduplication module*

Seven parameters have been estimated in order to fully calibrate the division-endoreduplication module (Bertin et al. 2007). Briefly, $\tau$ and $\tau_E$ represent the duration of the mitotic cell cycle and of the endocycle, respectively. The four parameters (theta_0, theta_M, mu and a) parameterize the Fermi function describing the decrease of cell proliferating activity with time ( $\theta(t)$ ), whereas $\sigma$ represents the proportion of expanding cells performing a new round of endoreduplication, after a time t $\tau_E$ (the minimum time required to complete an endocycle).

Calibration was performed using genetic algorithm (R language, 'genalg' package ), and repeated ten to twenty times in order to assure a good exploration of the parameter space. Parameters range have been established based on literature information and previous works (Bertin et al., 2007).

Data from the 2004 experiment on the evolution of cells number and ploidy levels during fruit development have been used at first. The dynamics of cell division (5 parameters) was then re-estimated on cell number measured in the 2007 experiment, whereas parameters $\tau_E$ and $\sigma$ (relative to endoreduplication dynamics) were kept fixed to values estimated on the 2004 data. The best-fitting parameters, for both studied genotypes, are reported in the following table:

| | | $\tau$ (hours) | $\tau_E$ (hours) | theta0 | thetaM | mu | a | $\sigma$ |
|---|---|---|---|---|---|---|---|---|
| **CERVIL** | **2004** | 18 | **13** | 0.66 | 0.39 | 15.53 | 7.5 | **0.046** |
| | ***2007*** | 28 | *NA* | 0.71 | 0.05 | 18.45 | 2.6 | *NA* |
| **LEVOVIL** | ***2004*** | 27 | **18** | 0.66 | 0.465 | 6.77 | 0.19 | **0.048** |
| | **2007** | 24 | *NA* | 0.60 | 0.45 | 18.06 | 3.53 | *NA* |

**Table S2**: Calibrated parameters values for the division-expansion module, for both genotypes, for the two experiments.

For both genotypes, estimated values for cell cycle duration are around 24 hours in agreement with literature information (Autran et al., 2002; Roeder et al., 2010). The entry into endoreduplication (endocycle) reduces the cell cycle duration in both genotypes.

The dynamics of cell division is strongly affected by the environment for the Cervil genotype. Cell cycle length is larger in 2007 than in 2004 (28h against 18h on average) but the duration of the division phase is shorter. Note that the initial cell value is also different between the two years, for both genotypes.

The time course of cell number and ploidy levels are reported in Fig. S4-S7 for the two genotypes and for the two experimental datasets.

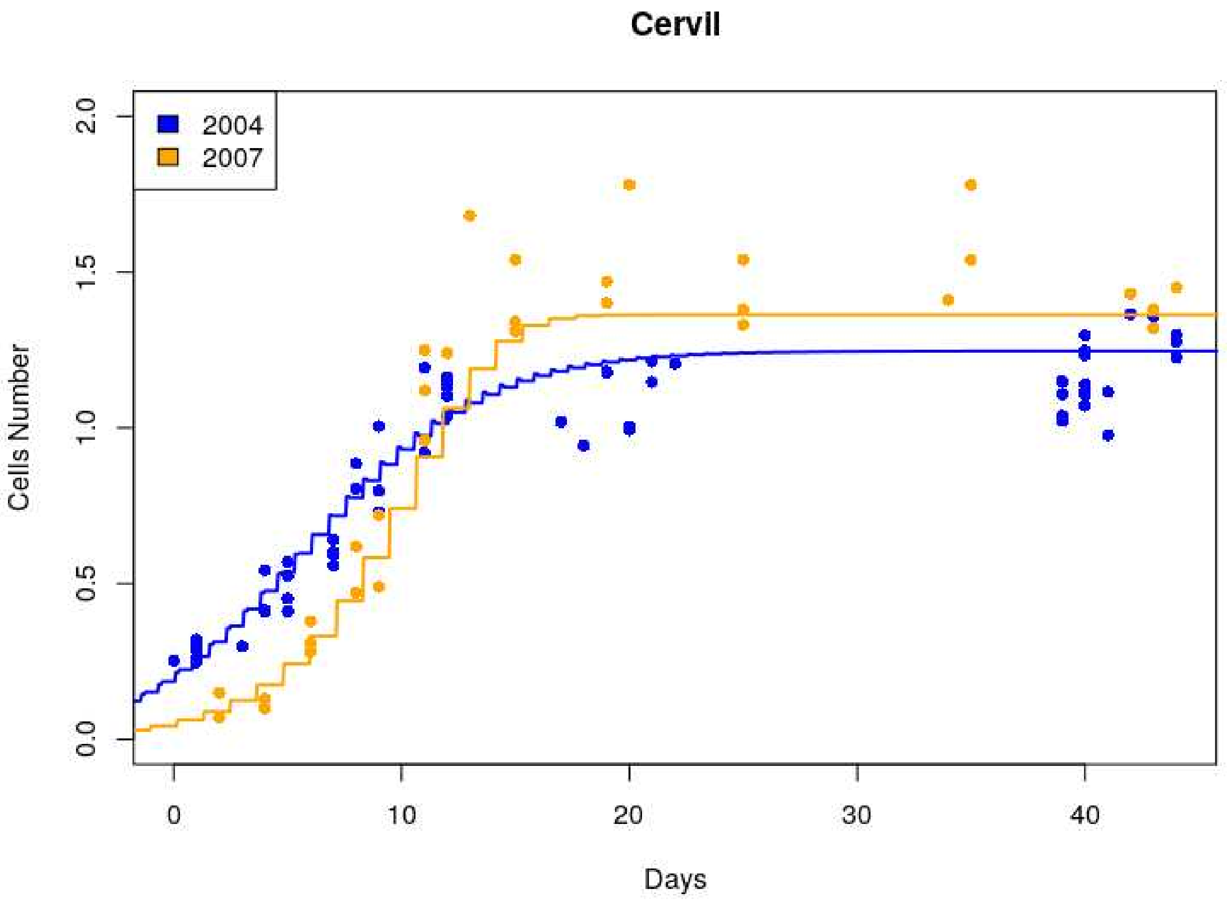


**Figure S3:** Fit of the dynamics of cell number for Cervil genotype. Dots represent data from the 2004 and 2007 experiment whereas the line represents the model simulation. Cell number is expressed in units of $10^6$ cells.

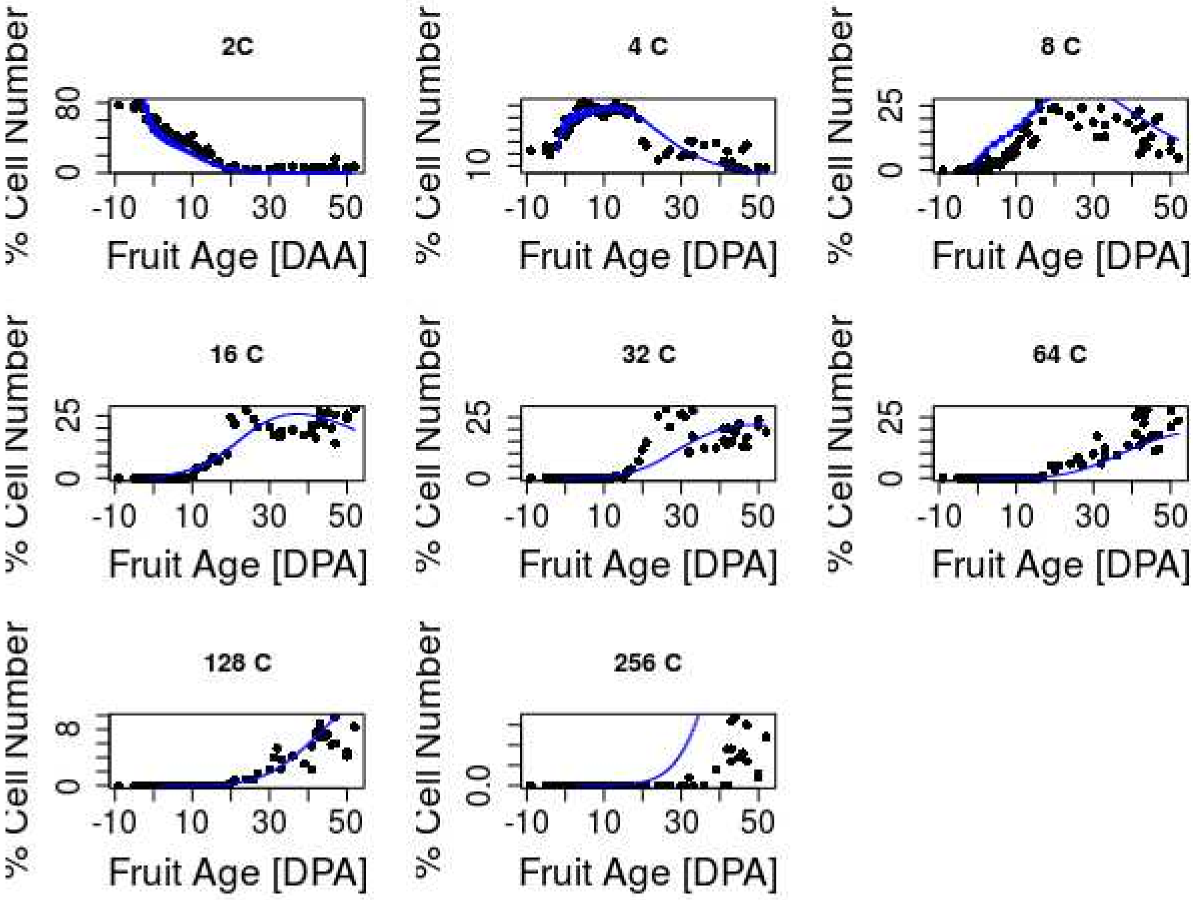


**Figure S4 :** Fit of the dynamics of endoreduplication for Cervil genotype. The temporal evolution of the fraction of cells in each ploidy class is shown. Dots represent data from the 2004 experiment whereas the line represents the model simulation.

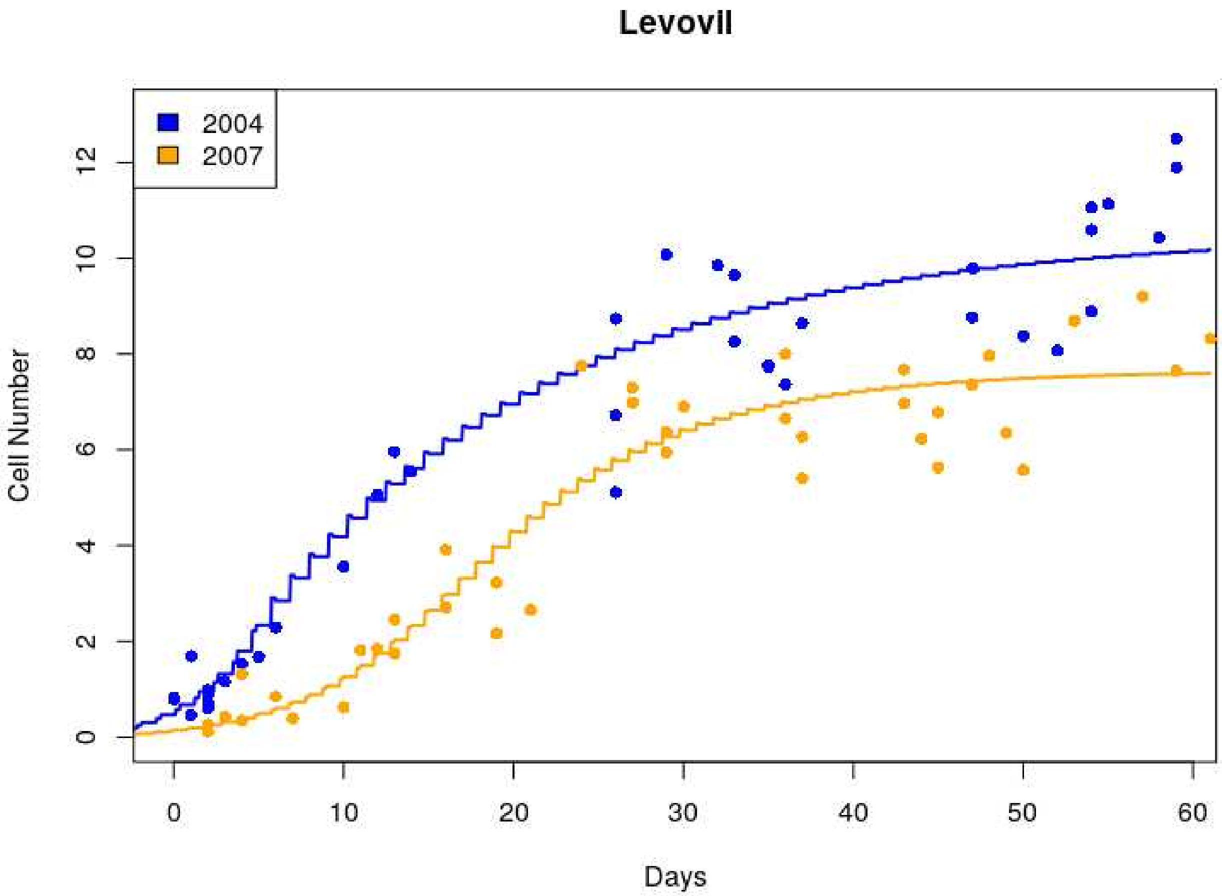


**Figure S5:** Fit of the dynamics of cell number for Levovil genotype. ots represent data from the 2004 and 2007 experiment whereas the line represents the model simulation. Cell number is expressed in units of $10^6$ cells.

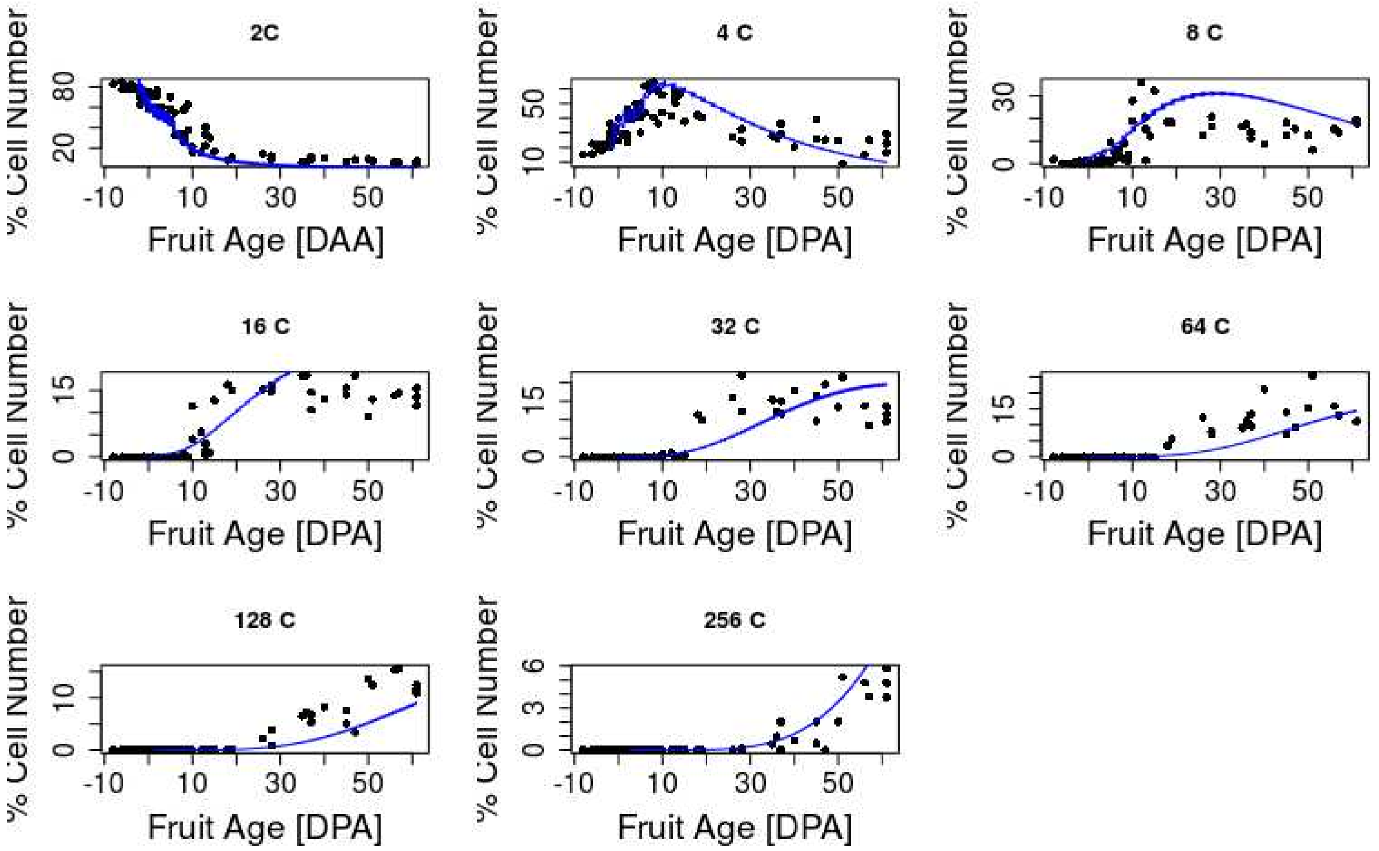


**Figure S6:** Fit of the dynamics of endoreduplication for Levovil genotype. The temporal evolution of the fraction of cells in each ploidy class is shown. Dots represent data from the 2004 experiment whereas the line represents the model simulation.

#### *S3.2 Calibration of the expansion module*

Ten variants of the expansion module have been tested and calibrated on the evolution of pericarp fresh and dry mass, from the 2007 experiment. 6 parameters have been selected for calibration based on a previous sensitivity analysis (Constantinescu et al., 2016) . Briefly, parameters $v_m, t_{star}, a_{ssrat}$ specify the dynamics of the active carbon uptake by the cell and its further use for the synthesis of soluble components. $\tau_S$ defines the timing for plasmodesmata closure during fruit development. $L_p$ and the ratio $L_x$ over $L_p$ specify the xylem and phloem conductivities.

If not indicated otherwise, the remaining parameters have been fixed to the original model values (Baldazzi et al., 2013; Fishman and Génard, 1998; Liu et al., 2007). However, in the case of a ploidy-dependent effect on cell soluble fraction (bssrat) or cell's plasticity (

$\phi_{max}$ ) we decided to re-calibrate the basal value of the corresponding processes in order to better evaluate the strength of the interaction between endoreduplication and expansion processes. For model version M3, M4, M6, M7, M23 and M24 therefore, seven parameters have been calibrated instead of six.

Calibration have been performed using genetic algorithm (R language, ‘genalg’ package ), and repeated three to five times for each model variant and genotype. The quality of model adjustment was evaluated using a Normalized Root Mean Square Error (NRMSE), as explained in the main text.

*3.2.1 Selection of the calibration solution for the expansion module*

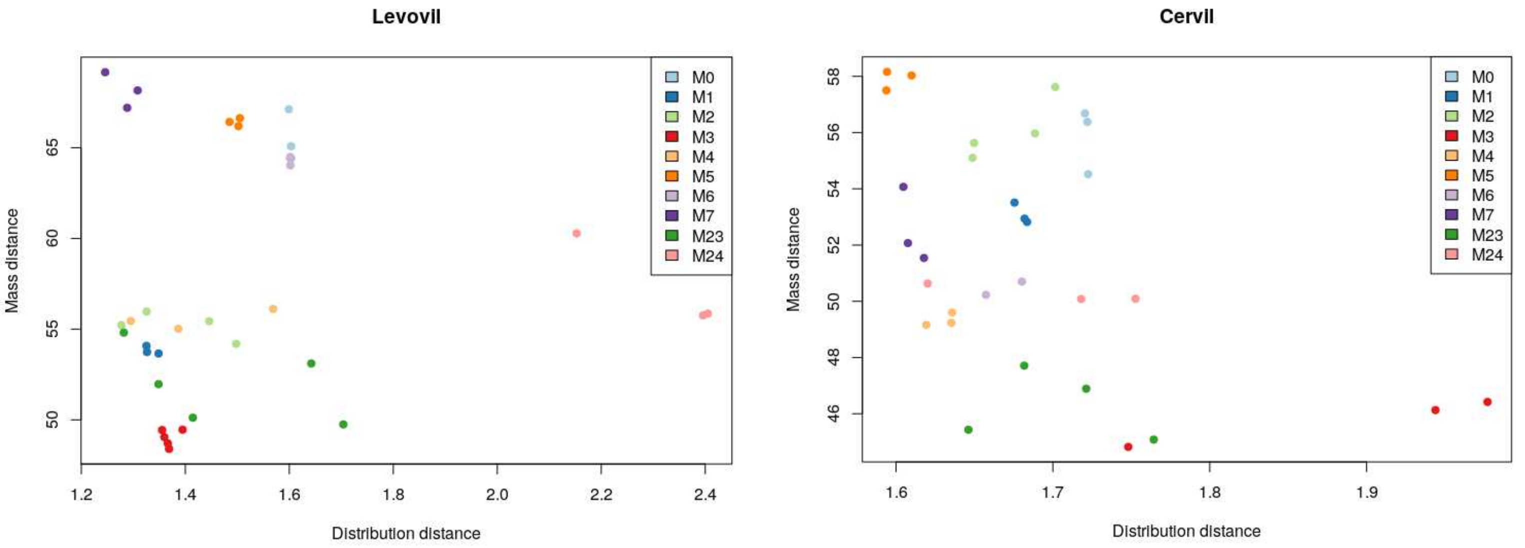


**Figure S7:** Repartition of all calibration solutions according to their agreement to observed pericarp mass (‘Mass distance’) and observed cell size distribution (‘Distribution distance’) at fruit maturity.

For each model version and calibration solution, we computed the cell size distribution predicted at fruit maturity and compared it to the experimental one. The calibration solution giving the best compromise between quality of the fit on the pericarp mass and quality of the distribution (in the sens of Pareto efficiency) was selected for each model version (see Figure S7). The corresponding estimated parameter values are reported in Table S3 and S4 for Cervil and Levovil genotype, respectively. A comparison between the simulated and the experimental dynamics of pericarp growth is plotted in figures S8 and S9.

| CERVIL | | | | | | | | |
|---|---|---|---|---|---|---|---|---|
| Model Variant | $v_m$ | $t_{star}$ | $a_{ssrat}$ | $\tau_S$ | Lx/Lp | Lp | $b^0_{ssrat}$ | $\phi_{max}$ |
| M0 | 0.19 | 15.73 | 0.99 | 9.9e-5 | 0.1 | 0.061 | NA | NA |
| M1 | 0.17 | 33.4 | 0.70 | 9.1e-6 | 0.19 | 0.055 | NA | NA |
| M2 | 0.075 | 9.6 | 0.70 | 1.0e-5 | 0.52 | 0.049 | NA | NA |
| M3 | 0.077 | 123.12 | 0.12 | 6.3e-6 | 0.10 | 0.51 | 0.016 | NA |
| M4 | 0.15 | 38.14 | 0.69 | 1.4e-5 | 0.10 | 0.41 | NA | 0.0015 |
| M5 | 0.065 | 17.14 | 0.69 | 5.6e-5 | 0.10 | 0.073 | NA | NA |
| M6 | 0.096 | 71.7 | 0.69 | 5.7e-5 | 0.35 | 0.27 | 0.014 | NA |
| M7 | 0.13 | 54.4 | 0.16 | 9.9e-5 | 0.10 | 0.50 | NA | 0.0022 |
| M23 | 0.051 | 52.0 | 0.10 | 1.04e-5 | 0.59 | 0.55 | 0.021 | NA |
| M24 | 0.049 | 51.6 | 0.70 | 6.5e-6 | 0.10 | 0.35 | NA | 0.0021 |

**Table S3:** Retained parameters values for the expansion module, model versions M0 to M24, for Cervil genotype.

| LEVOVIL | | | | | | | | |
|---|---|---|---|---|---|---|---|---|
| Model Variant | $v_m$ | $t_{star}$ | $a_{ssrat}$ | $\tau_S$ | Lx/Lp | Lp | $b^0_{ssrat}$ | $\phi_{max}$ |
| M0 | 0.017 | 500.5 | 0.50 | 6.8e-6 | 0.13 | 0.042 | NA | NA |
| M1 | 0.024 | 519.6 | 0.35 | 3.77e-6 | 0.20 | 0.047 | NA | NA |
| M2 | 0.006 | 492.8 | 0.38 | 2.8e-6 | 0.12 | 0.055 | NA | NA |
| M3 | 0.020 | 569.4 | 0.33 | 4.3e-6 | 0.23 | 0.12 | 0.016 | NA |
| M4 | 0.032 | 403.3 | 0.31 | 3.9e-6 | 0.10 | 0.051 | NA | 0.003 |
| M5 | 0.003 | 433.5 | 0.16 | 2.2e-6 | 0.10 | 0.046 | NA | NA |
| M6 | 0.018 | 400.4 | 0.50 | 4.2e-6 | 0.19 | 0.037 | 0.0008 | NA |
| M7 | 0.026 | 404.4 | 0.17 | 7.3e-6 | 0.10 | 0.17 | NA | 0.002 |
| M23 | 0.006 | 493.0 | 0.59 | 1.9e-6 | 0.41 | 0.059 | 0.07 | NA |
| M24 | 0.010 | 402.1 | 0.17 | 8.8e-6 | 0.10 | 0.24 | NA | 0.002 |

**Table S4:** Retained parameters values for the expansion module. Model versions M0 to M24, for Levovil genotype.

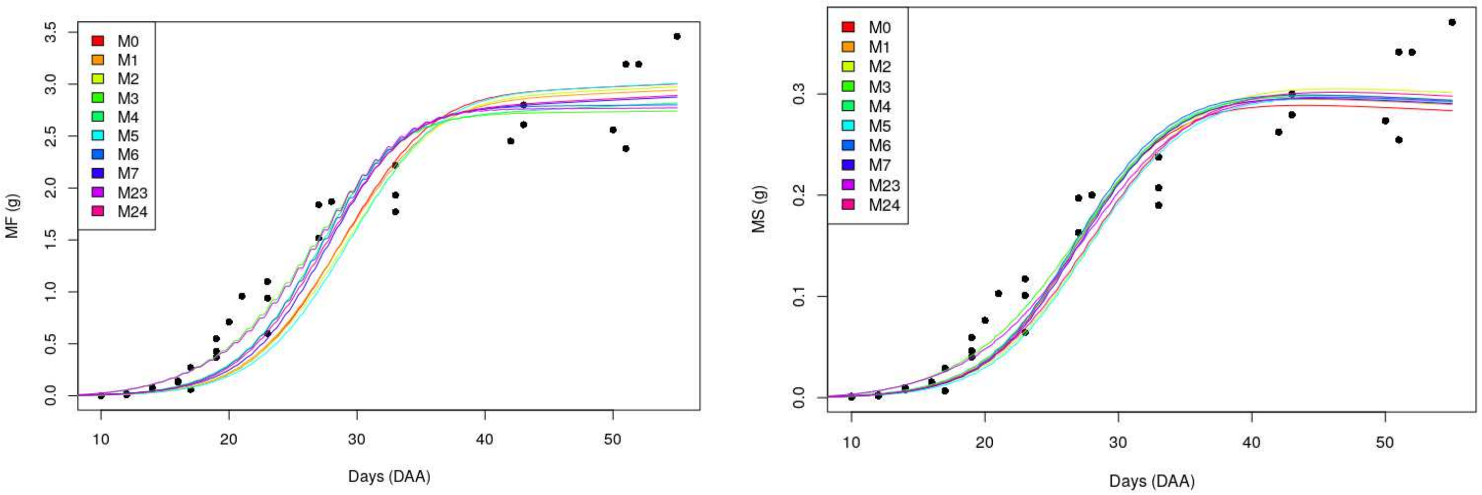


**Figure S8:** Calibration of pericarp growth with models M0-M24 for Cervil genotype. Points represent experimental data, lines represent the selected model simulations. Left: Pericarp fresh mass. Right: Pericarp dry mass.

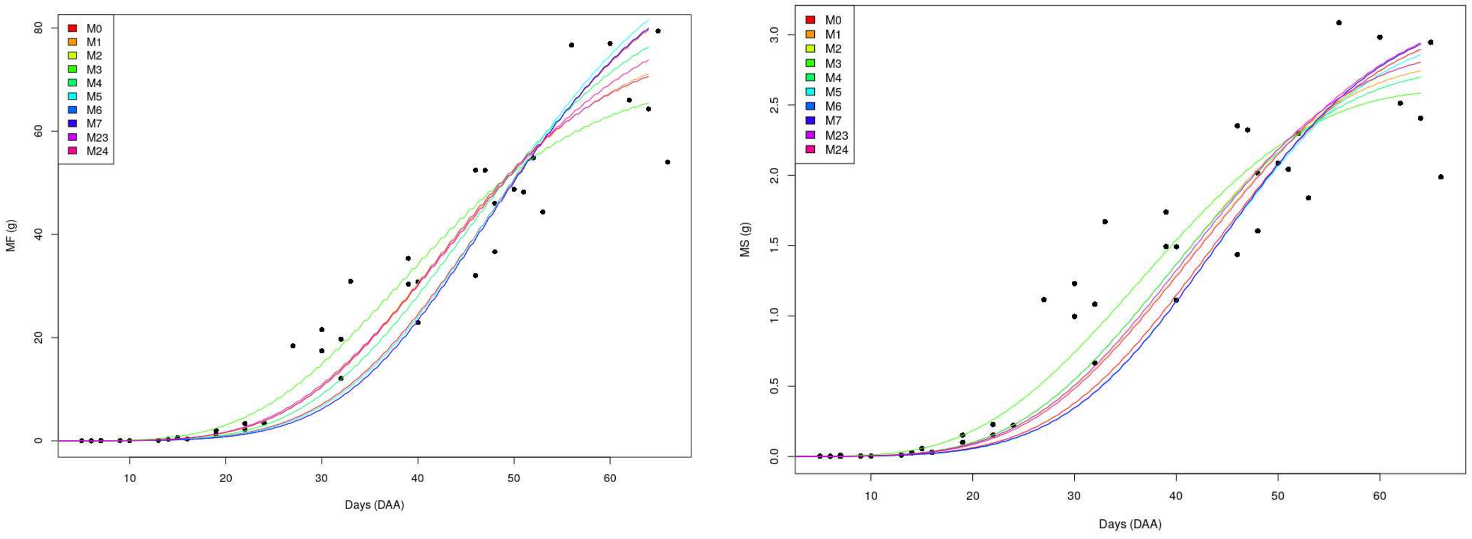


**Figure S9:** Calibration of pericarp growth with models M0-M24 for Levovil genotype. Points represent experimental data, lines represented the selected model simulations. *Left:* Pericarp fresh mass. *Right:* Pericarp dry mass.

**S4: Correlation between ploidy and size**

Correlation between ploidy and size was systematically tested for all models. To this aim, a linear regression analysis was performed using R software. Hereafter we report the estimated coefficients of the regression model :

cell area (mm2) ~ y0 +m* ploidy

over all fruit cells at maturity.

Model adjusted-R2 and significance of the *m* term (p-value) are also reported.

| | **CERVIL** | | | | **LEVOVIL** | | | |
|---|---|---|---|---|---|---|---|---|
| Model | y0 | m | | R2 | y0 | m | | R2 |
| M0 | 2.7e-2 | -3.5e-7 | *** | 0.007 | 7.3e-2 | 3.7e-5 | *** | 0.04 |
| M1 | 2.7e-2 | 2.1e-6 | *** | 0.024 | 6.1e-2 | 1.0e-4 | *** | 0.08 |
| M2 | 1.8e-2 | 9.2e-5 | *** | 0.88 | 4.7e-2 | 2.3e-4 | | 0.21 |
| M3 | 2.1e-2 | 2e-5 | *** | 0.056 | 5.3e-2 | 1.2e-4 | *** | 0.09 |
| M4 | 2.2e-2 | 2.2e-5 | *** | 0.65 | 5.6e-2 | 1.9e-4 | *** | 0.28 |
| M5 | 2.0e-2 | 5.9e-5 | *** | 0.82 | 6.9e-2 | 5.6e-5 | *** | 0.09 |
| M6 | 2.2e-2 | 6.4e-6 | *** | 0.49 | 6.8e-2 | 6e-5 | *** | 0.06 |
| M7 | 2.2e-2 | 2.4e-5 | *** | 0.70 | 5.4e-2 | 2.9e-4 | *** | 0.63 |
| M23 | 1.8e-2 | 7.8e-5 | *** | 0.54 | 5.0e-2 | 2.5e-4 | *** | 0.41 |
| M24 | 1.8e-2 | 9.2e-5 | *** | 0.88 | 2.7e-2 | 6e-4 | *** | 0.64 |

**Table S5:** Estimated parameters for the linear regression model testing the hypothesis of a linear relation between cell size (mm2) and its ploidy level. Significance is indicated as follows: *** p-value <0.001, ** pvalue <0.01 * pvalue<0.05, '.' pvalue <0.1, '-' elsewhere)

In order to better investigate the origins of correlation between size and ploidy, maximal growth rate was computed as

$$max_t \frac{dFW_i}{dt}$$

where *i is t*he ploidy class and *t* is cell developmental time (cell age)
Correlation between ploidy and maximal growth rate was then quantified by means of the Pearson correlation coefficient (table S6). Function cor.test of R software has been used to

this purpose. Significance of the result is reported following the code: *** p-value <0.001, ** p-value <0.01, * p-value<0.05, '.' pvalue <0.1, '-' elsewhere.

| Model | CERVIL | | LEVOVIL | |
|---|---|---|---|---|
| | Pearson Correlation | Significance | Pearson Correlation | Significance |
| M0 | 3e-7 | - | 0.07 | - |
| M1 | 7e-16 | - | 0.03 | - |
| M2 | 0.85 | *** | 0.31 | *** |
| M3 | 0.081 | - | 0.077 | . |
| M4 | 0.67 | *** | 0.29 | *** |
| M5 | 0.90 | *** | 0.30 | *** |
| M6 | 0.84 | *** | 0.054 | - |
| M7 | 0.78 | *** | 0.87 | *** |
| M23 | 0.79 | *** | 0.42 | *** |
| M24 | 0.85 | *** | 0.37 | *** |

**Table S6:** Pearson correlation coefficient between the maximal cell growth rate and the ploidy level. Significance is indicated as follows: *** p-value <0.001, ** p-value <0.01, * p-value<0.05, '.' pvalue <0.1, '-' elsewhere)
